# Implication of *TITIN* Variations in Dilated Cardiomyopathy: Integrating Whole Exome Sequencing With Molecular Dynamics Simulation Study

**DOI:** 10.1101/2024.11.10.622829

**Authors:** Amrita Mukhopadhyay, Bharti Devi, Anurag TK Baidya, Manohar Lal Yadav, Rajnish Kumar, Bhagyalaxmi Mohapatra

## Abstract

Dilated cardiomyopathy (DCM) is one of the leading causes of heart failure, characterized by ventricular dilation and impaired systolic function. Variations in the *TITIN* (*TTN)* gene, which encodes the giant muscle protein TTN, play a pivotal role in the genetic underpinnings of DCM. We conducted WES on 15 patients (5 familial and 10 sporadic) diagnosed with idiopathic DCM and identified 88 exonic variants including four novel variants. These variants were predominantly located in the A-band region (39 variants) of TTN, a critical region for its mechanical stability and interaction with other sarcomeric proteins, followed by the I-band domain (33 variants), Z-disc domain (7 variants) and M-band region (9 variants). To discern the functional repercussions of these variations, we performed several bioinformatics analyses including pathogenicity prediction, protein stability, and protein-protein docking followed by MD simulations on both wild-type and mutant TTN fragments with their corresponding interacting partners (TCAP, MYH7, LMNA). We revealed that variations in the A-band domain significantly alter the protein’s structural dynamics, leading to decreased mechanical stability and altered protein-protein interactions. These changes are likely to disrupt sarcomere function, thereby elucidating their role in the pathogenesis of DCM.

**Graphical abstract:** 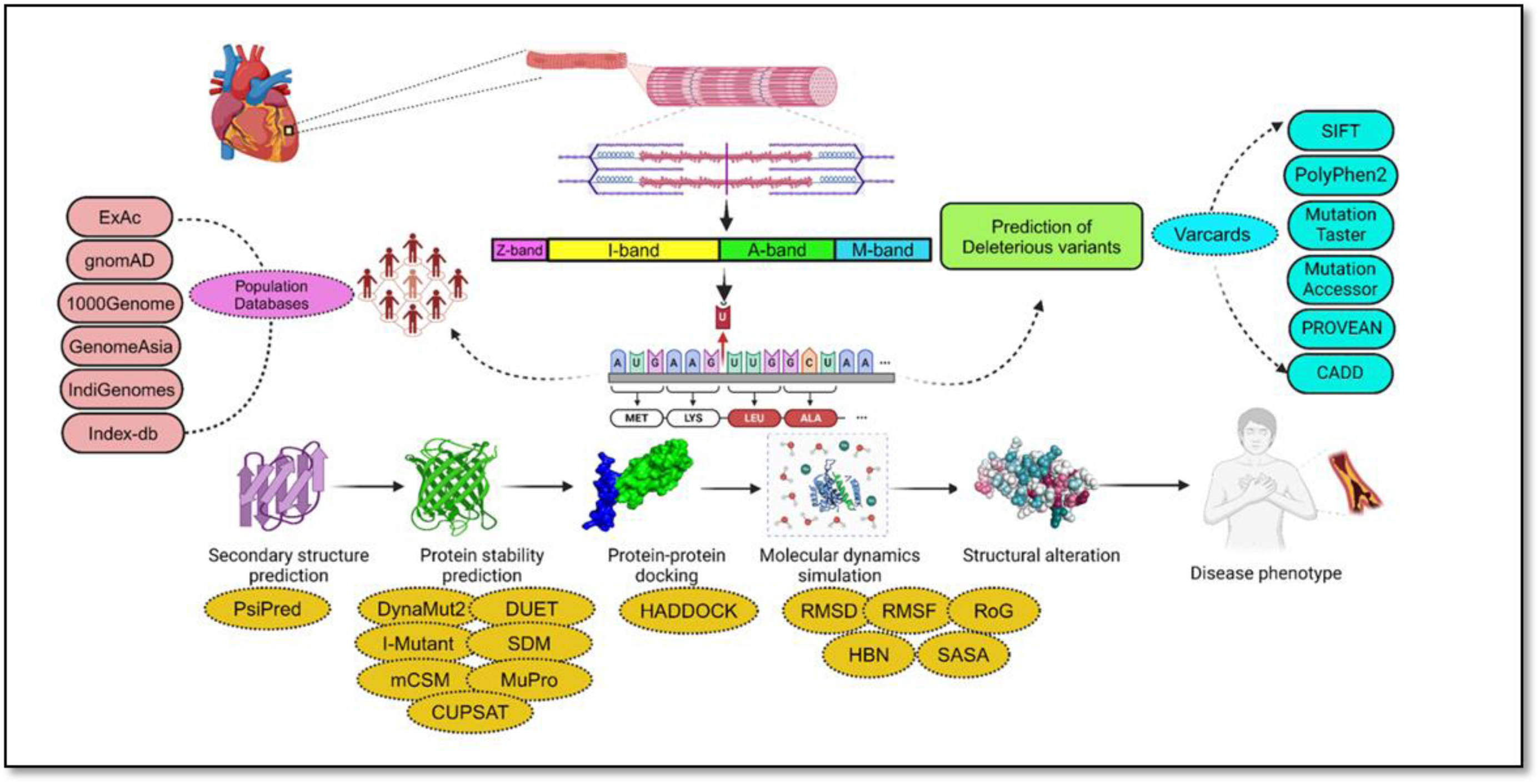

## 1 Introduction

DCM represents the most prevalent form of cardiomyopathy, affecting approximately 1 in 2500 individuals [1]. It is characterized by dilation of left- or bi-ventricles and impaired systolic function that ultimately leads to progressive cardiac failure [2]. DCM of unknown cause is clinically classified as idiopathic DCM and 20-35% of idiopathic DCM cases are attributed to genetic transmission [3]. More than 60 genes have been reported to be associated with DCM thus far, most of which are involved in force generation and force transmission, involving sarcomeric and cytoskeletal genes [4]. Among them, *TTN* plays a pivotal role, contributing substantially to the development of DCM.

TTN, the gigantic human protein (∼4 MDa), is considered a key component for sarcomere integrity and function [5, 6]. The enormous size (294 kb) and complexity of the *TTN* hampered variational analysis of the gene by conventional Sanger sequencing previously [7]. The advent of NGS technologies has enabled the diagnosis of the spectrum of variations in the *TTN* gene. Herman et al. (2012) were the first to report variations in the *TTN* using NGS technology, by analyzing 312 patients with DCM and 231 patients with hypertrophic cardiomyopathy (HCM) [8]. They detected *TTN* truncating variations in 25% of familial DCM and in 18% of sporadic DCM cases. However, reports of functional analysis of the identified variations are scanty.

Due to the unavailability of a commercial clone for the full-length *TTN* and the challenges of conducting *in-vitro* and *in-vivo* experiments with the entire protein, we opted for *in-silico* approaches to analyse the variants identified through Whole Exome Sequencing (WES) in 15 patients (10 sporadic cases and 5 familial cases) with DCM. We identified a total of 88 coding variants, of which 21 variants were classified as ‘damaging’. Out of these 21 variants, we prioritized 7 variants based on their novelty and possible functional impact. We used several bioinformatics algorithms to determine secondary-structure, folding pattern, domain functions, protein stability and conservation analysis of native and mutant protein complexes. By using Alpha-fold2, we constructed 3D models of TTN and its associated interacting partners followed by protein-protein docking studies to elucidate the interactions between TTN and other protein molecules. The reliability of the docking results was evaluated through comprehensive Molecular Dynamics (MD) simulations. MD simulations revealed significant structural changes in mutant forms, indicating potential impairment of TTN function in DCM pathogenesis. Thus our study put a spotlight on *TTN* variations identified through WES in familial and sporadic cases of DCM implying that these variations could be causative factors in the pathogenesis of DCM.

## 2 Materials and method

### 2.1 Enrolment of study subjects and clinical evaluation

A total of fifteen idiopathic dilated cardiomyopathy patients (age<25years), including 5 familial (FDCM1-5) and 10 sporadic cases (SDCM1-10) were recruited for this study from the Department of Cardiology and the Department of Pediatric Medicine, SS Hospital, Institute of Medical Sciences, Banaras Hindu University after written informed consent. Family studies include at least two affected individuals within the same family. Diagnosis relied on the LV dimensions and LV functions (i.e, LV ejection fraction (LVEF) of <45% and fractional shortening (LVFS) <25%)). Patients with hypertension, coronary artery disease (CAD) and congenital heart disease (CHD) were excluded from the study. Study approval was obtained from the Institutional Ethics Committee.

### 2.2 Genomic DNA isolation and Library preparation for Whole Exome Sequencing

Genomic DNA (gDNA) was extracted from the collected blood samples by standard ethanol precipitation method. The gDNA library was prepared using human Nextera Rapid Capture Expanded Exome kit (Illumina, USA) according to the manufacturer’s instructions. Prepared libraries were sequenced on the Illumina HiSeq2500 platform (Illumina HiSeq2500, USA) to generate 2×150bp paired-end reads per sample with a coverage of 100x. The produced sequence data underwent the required quality control checks.

### 2.3 NGS data analysis

Data Analysis was performed following the completion of sequencing. After aligning the data to the human reference genome (GRCh38: Genome Reference Consortium Human Build 38 or hg19), PCR duplicates were removed by Picard-tools-2.1.1. Reads were re-aligned around the known indels using GATK (Genome Analysis ToolKit), recalibrated using GATK Toolkit-Recalibrator and VCF files were generated for each sample. Details methodology of NGS data analysis is available on a request basis.

### 2.4 Variant prioritization

The potentially disease-causing variants were prioritized using the pipeline outlined (**Figure 1**). In summary, the variants were initially selected having read-depth ≥ 20 and then matched with our curated gene panel. Variants were further filtered based on a minor allele frequency (MAF) of <0.01 in the Genome Aggregation Database (gnomAD; https://gnomad.broadinstitute.org/), the 1000 Genomes Project (https://www.internationalgenome.org), and the ExAC database (https://exac.broadinstitute.org/). The selected variants were categorized separately into missense, frame-shift (insertion/deletion), nonsense (stop-codon), synonymous (silent), splice-site and intronic variants, depending on their impact on the amino acid sequence.

**Figure 1:**
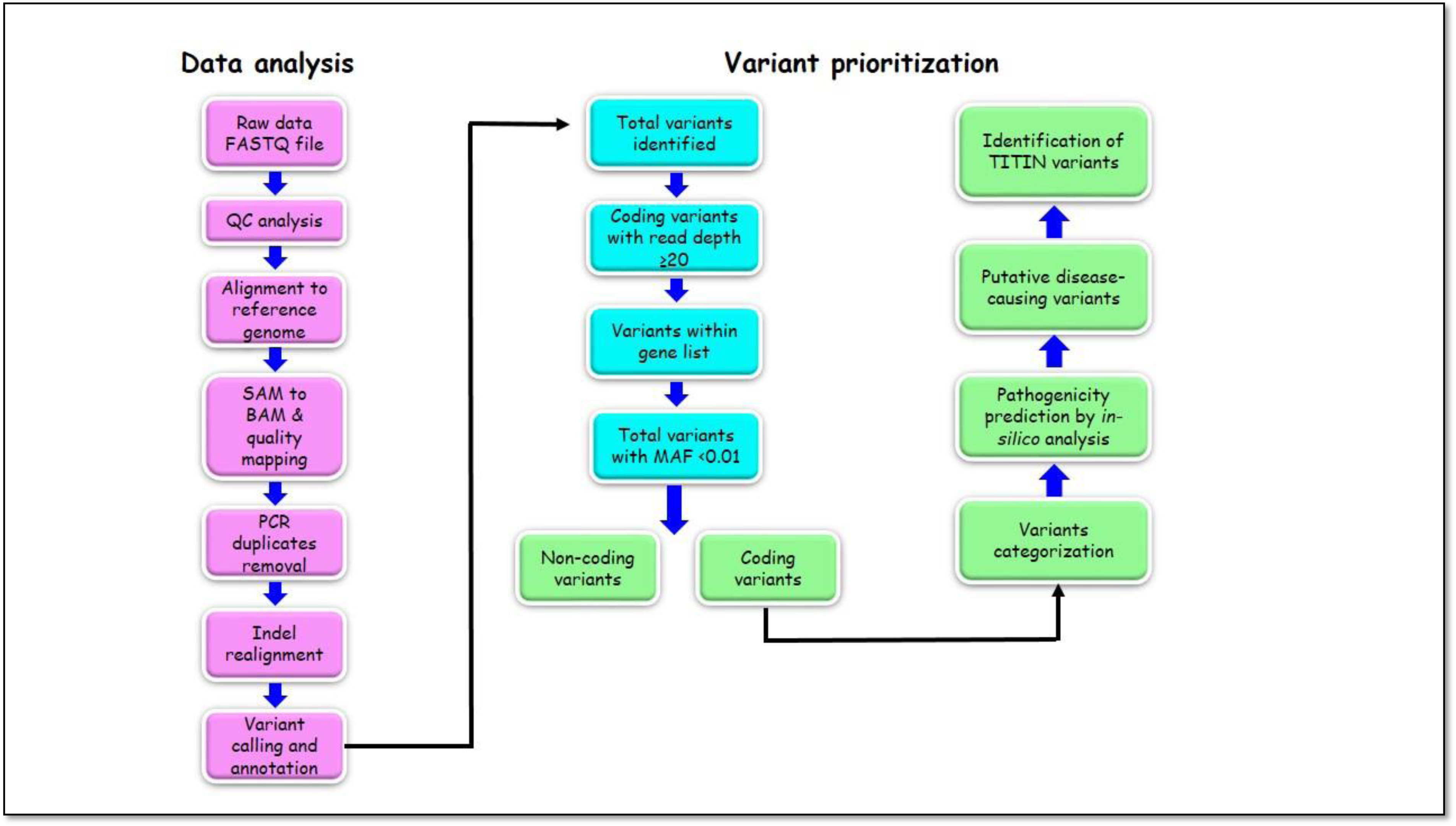
WES pipeline for data analysis and variant prioritization.

### 2.5 Pathogenicity prediction of selected variants through *in-silico* tools

An *in-silico* analysis was conducted using Varcards (http://www.genemed.tech/varcards/search) to determine the pathogenic potential of coding variants [9]. The study incorporated over 20 tools (https://brb.nci.nih.gov/seqtools/colexpanno.html) to predict the damaging scores (ranging from 0 to 1) of coding variants at the protein structure and function levels. Some of the tools utilized in the analysis included SIFT, PolyPhen2_HDIV, PolyPhen2_HVAR, FATHMM, VariationTaster, VariationAssessor, PROVEAN, MetaSVM, MetaLR, M-CAP, CADD, VEST3, GERP++, GenoCanyon, DANN, fitCons, PhyloP, SiPhy, and REVEL.

### 2.6 Phylogenetic conservation of the damaging variants and secondary structure prediction

To assess the conservation of substituted amino acids for homologs across different species, the COBALT at the NCBI (https://www.ncbi.nlm.nih.gov/tools/cobalt/re_cobalt.cgi) was used. Additionally, the Consurf (https://consurf.tau.ac.il/consurf_index.php) tool [10] was used to determine the localization of the residues whether exposed or buried in the protein domains. Further, we compared the effects of variations on the secondary structures of wild-type (WT) and mutant proteins using the PsiPred server (http://bioinf.cs.ucl.ac.uk/psipred/) [11].

### 2.7 Protein modelling and structure validation

Unfortunately, the crystal structure of full-length TTN and other interacting partners has not yet been resolved. Thus the sequences in FASTA format were retrieved from Universal Protein Resource ‘UniProt Database’ (Uniprot ID: TTN-Q8WZ2, ACTC1-P68032, MYH7-P12883, LMNA-P02545). The 3D structure of TCAP was obtained from RCSB PDB (TCAP: 1YA5). Then the sequences were submitted to AlphaFold2 googlecolab (https://www.nature.com/articles/s41586-021-03819-2) for the prediction of 3D model of the protein. Thenceforth, the structural and energetic validation was performed using the molecular dynamic (MD) simulation for the modeled structures for 10ns.

### 2.8 Protein-Protein docking

The refined structure that we got after the MD simulation was used to check molecular interactions. To reveal the interaction of TTN with other protein molecules, we performed molecular docking studies using HADDOCK2.4 (https://doi.org/10.1016/j.jmb.2015.09.014) webserver [12]. PDB structures of the docking partners were uploaded on the webserver and binding residues were also provided for each of the partners based on literature survey [13–23]. For mutated structures, the regions including variation were selected as active/binding residues that were directly involved in the interaction. Structures were visualized with the help of PyMOL 4.6 software.

### 2.9 Molecular Dynamics Simulation

The complex/cluster that demonstrated high binding affinity was selected for carrying out classical molecular dynamic simulation for 200 ns using software GROMACS version 2020 [24]. Initially, to construct the system, we used CHARMM-GUI web server [25] employing CHARMM 36m force field [26,27]. For solvation of the protein, TIP3P water molecules [27] were utilized in a cubic box size of appropriate dimensions and neutralized with Na^+^ and Cl^-^ ions by means of Monte Carlo ion placing method. Energy minimization comprised 50,000 steps with steepest descent algorithm until forces were below 1000 (kJ/mol)/nm, , followed by 2 ns of NVT (constant number of particles, Volume and Temperature) ensemble equilibration with protein restraints at 310.15 K using a Noose Hoover thermostat with 1.0 ps coupling constant for all atoms. Subsequently, NPT ensemble equilibration continued for 2 ns at 1 bar pressure using a Parrinello-Rahman barostat [28] with a 1 ps coupling constant. Finally, the production simulation extended for 200 ns under isobaric-isothermal conditions, employing a time step of 2 fs. H-bond constraints were managed by the LINCS algorithm [29] and long-range electrostatic interactions were treated using the Particle mesh Ewald (PME) method, with a 1.2nm cut-off and 0.16 nm grid spacing. Van der Waals interactions were truncated at 1.2 nm. Trajectories were analyzed post-simulation for root mean square deviation (RMSD), root mean square fluctuations (RMSF), radius of gyration (RoG), hydrogen bond number (HBN), hydrogen bond distance (HBD), solvent accessible surface area (SASA), principal component analysis (PCA) and free energy landscape (FEL) analysis with the help of GROMACS.

### 2.10 Protein stability prediction by miscellaneous tools

Different approaches namely, Dynamut2 (https://biosig.lab.uq.edu.au/dynamut2/) [30], MUpro (https://mupro.proteomics.ics.uci.edu/) [31], CUPSAT (https://cupsat.brenda-enzymes.org/) [32], DUET (https://biosig.lab.uq.edu.au/duet/stability [33], I-mutant (https://folding.biofold.org/i-mutant/i-mutant2.0.html) [34] were used to assess changes in protein flexibility and stability upon variations. To study physicochemical properties of mutant structures, ProtParam (https://web.expasy.org/protparam/) [35] tool was used. Further, PDbsum tool (http://www.ebi.ac.uk/thornton-srv/databases/pdbsum/) [36] was used to generate a Ramachandran plot as well as 2D representation of intermolecular interactions for each complex.

### 2.11 Statistical analysis

Pearson’s χ² (chi-squared) goodness-of-fit tests were employed to examine the random distribution of the disease-causing variants. The analysis incorporated two groups; taking the size in amino acids of each protein region tested as one group and the remainder of TTN protein as a second group (df=1). This statistical approach allowed us to assess whether the distribution of variants across these two groups followed an expected pattern or deviated significantly from randomness [37].

## 3. Results

### 3.1 Identification of *TTN* variants

Whole exome sequencing of five families and 10 sporadic DCM cases revealed a total of 92 genetic variations (88 exonic, 3 intronic, and 1 UTR) in the *TTN* gene (**Table 1**, **Supplementary Table 1**). Of the 88 exonic variants, 68 were non-synonymous, 17 were synonymous, 2 were frame-shift deletions, and 1 nonsense variation. Among these, one non-synonymous (c.56525G>A, R18842H), two frameshift-deletions (c.44969AG>A, p.A14990GfsTer4 and c.78090_78094delCTTTCT>C, p.E24390KfsTer15) and one nonsense (c.101629C>T, Q33877X) variations were identified as novel. All the novel variants were submitted to the Clinical Variation database with the following submission IDs: c.101629C>T (SCV002567998), c.44969delinsA (SCV3836535), c.56525G>A (SCV004015148), c.78090delinsC (SCV004015149). These four novel variants were not reported in gnomAD, 1000Genome and ExAC databases. Furthermore, none of these variants were found neither in GenomAsia database nor in any of the Indian databases, namely IndiGenom and Index-db. Out of 88 coding variants, we categorized 21 variants as ‘damaging’ based on the Varcards score. The list of allele frequency of these 21 damaging variants in different population databases, such as ExAc, gnomAD, and 1000Genome, as well as in Index-db, IndiGenom, and GenomAsia was shown in **Supplementary Table 2**. The minor allele frequency of all these variants in both the Indian databases were reported to be below 0.1%.

**Table 1:** List of damaging variants in TTN and their pathogenicity identified in DCM patients.

| Nucleotide change<br>(Amino acid change) | dbSNP | Varcards<br>Damaging<br>score | TTN domains | Interacting<br>partners |
| --- | --- | --- | --- | --- |
| c.178G>T<br>(D60Y) | rs35683768 | 0.67 | Z-disc | Telethonin |
| c.44969AG>A<br>(A14990GfsTer4) | Novel (SCV3836535) | -- | FnIII 8 (A-band) | Myosin |
| c.56525G>A<br>(R18842H) | Novel (SCV00401518) | 0.50 | Ig-like 107 (A-band) | Mysoin |
| c.67664G>A<br>(R22555H) | rs200317412 | 0.86 | FnIII 63 (A-band) | Mysoin |
| c.78090_78094del<br>CTTTCT>C<br>(E24390Kfster15) | Novel (SCV00401519) | -- | FnIII 77 (A-band) | Mysoin |
| c.90358G>A<br>(E30120K) | rs727505342 | 0.68 | FnIII 119 (A-band) | Mysoin |
| c.101629C>T<br>(Q33877X) | Novel (SCV00256798) | 1.0 | Ig-like 149 (M-band) | Lamin |

Abbreviations: rs-ID, Reference SNP cluster ID; TTN, Titin; Ig-like, Immunoglobulin-like, FnIII-Fibronectin type III;SCV, SCV, Clinical Variantion database accession numbers of the submitted novel variants

### 3.2 Distribution of *TTN* variants across protein domains

In our study cohort, we made a noteworthy discovery of 88 variations spanning various domains of *TTN*. When mapped to their corresponding protein domains, it was noted that 7 non-synonymous variants were localized to the Z-disc binding region of *TTN,* while 33 variants were confined to the I-band region followed by 39 variants localized at the A-band region, which include 2 frame-shift deletions. Following these, the M-band region also harboured 9 variations. Therefore, a significantly higher number of variations were residing at the Myosin-binding A-band region of TTN. We observed distribution of the variants followed non-random association (p-value =0.008 which is <0.05), i.e., the location of variations is biased towards the A-band region. Further, 3 out of the 4 novel variations found in our cohort were present in the Myosin-binding A-band region; the non-synonymous variation (R18842H) was present at Ig-like107 domain followed by two frame-shift deletions A14990GfsTer4 and E24390KfsTer15 at FnIII 8 and FnIII77 domains, respectively. Besides these, the novel stop-codon variation (Q33877X) was found to be located at the M-band region. Two other deleterious variations (c.30913G>A, p.G10305R and c.35183C>G, p.P11728R) were present at PEVK rich spring-like domain (I-band region) that altered the stiffness of the TTN by protein kinase phosphorylation (**Figure 2**).

**Figure 2:**
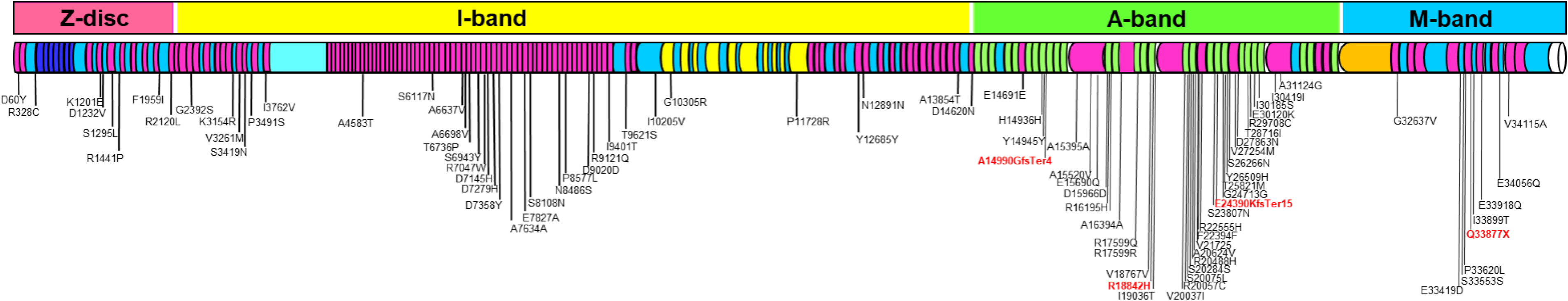
Distribution of TTN variants across protein domains.

### 3.3 Pathogenic potential discloses a significant number of potentially disease-causing *TTN* variants

The Varcards score ranges from 0 to 1, with higher scores indicating a higher likelihood of the variant being pathogenic. Based on the predictions, we chose 7 ‘damaging’ variants (D60Y, R29708C, R22555H, A14990GfsTer4, R16195H, E15690Q, A31124G) from familial cases and 14 variations (R20488H, V27254M, S20075L, A15520V, S23807N, E24390KfsTer15, R18842H, R20057C, P11728R, Y26059H, E30120K, Q33877X, I19036T, T25821M) in sporadic cases. The Varcards prediction scores of the damaging variants are shown in **Table 2**.

**Table 2:**
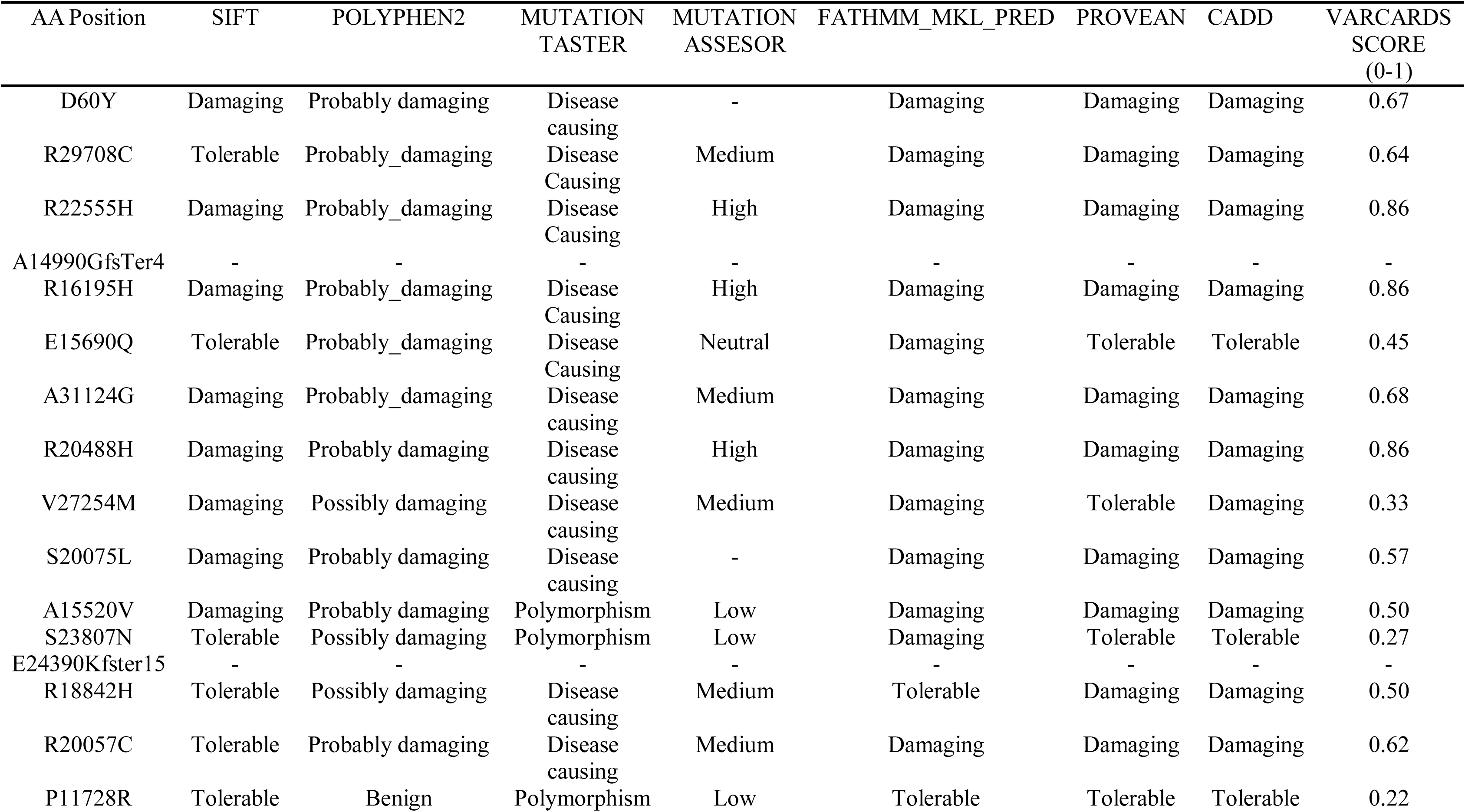
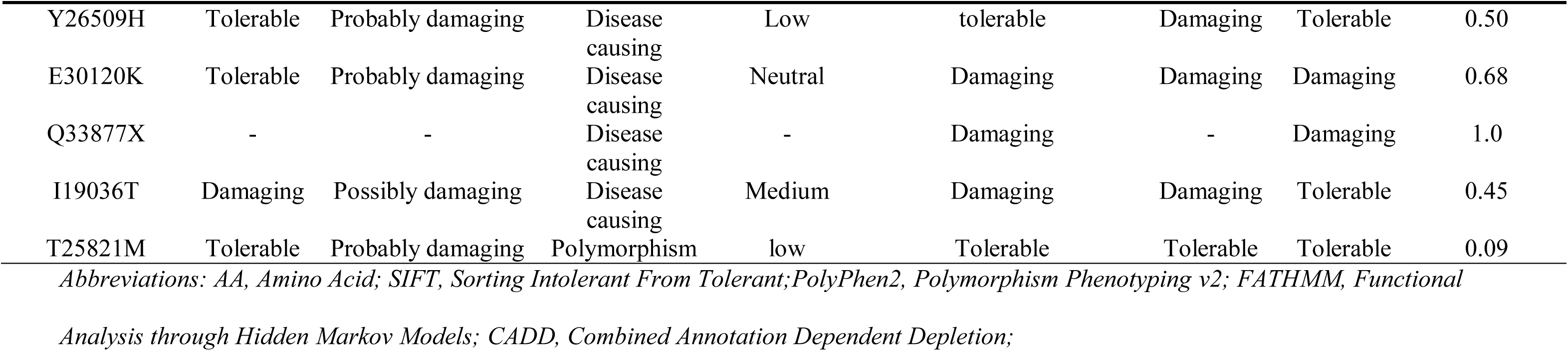
Prediction of the pathogenic likelihood of TTN variants using VarCards.

| AA Position | SIFT | POLYPHEN2 | MUTATION<br>TASTER | MUTATION<br>ASSESOR | FATHMM_MKL_PRED | PROVEAN | CADD | VARCARDS<br>SCORE<br>(0-1) |
| --- | --- | --- | --- | --- | --- | --- | --- | --- |
| D60Y | Damaging | Probably damaging | Disease causing | - | Damaging | Damaging | Damaging | 0.67 |
| R29708C | Tolerable | Probably_damaging | Disease Causing | Medium | Damaging | Damaging | Damaging | 0.64 |
| R22555H | Damaging | Probably_damaging | Disease Causing | High | Damaging | Damaging | Damaging | 0.86 |
| A14990GfsTer4 | - | - | - | - | - | - | - | - |
| R16195H | Damaging | Probably_damaging | Disease Causing | High | Damaging | Damaging | Damaging | 0.86 |
| E15690Q | Tolerable | Probably_damaging | Disease Causing | Neutral | Damaging | Tolerable | Tolerable | 0.45 |
| A31124G | Damaging | Probably_damaging | Disease causing | Medium | Damaging | Damaging | Damaging | 0.68 |
| R20488H | Damaging | Probably damaging | Disease causing | High | Damaging | Damaging | Damaging | 0.86 |
| V27254M | Damaging | Possibly damaging | Disease causing | Medium | Damaging | Tolerable | Damaging | 0.33 |
| S20075L | Damaging | Probably damaging | Disease causing | - | Damaging | Damaging | Damaging | 0.57 |
| A15520V | Damaging | Probably damaging | Polymorphism | Low | Damaging | Damaging | Damaging | 0.50 |
| S23807N | Tolerable | Possibly damaging | Polymorphism | Low | Damaging | Tolerable | Tolerable | 0.27 |
| E24390Kfster15 | - | - | - | - | - | - | - | - |
| R18842H | Tolerable | Possibly damaging | Disease causing | Medium | Tolerable | Damaging | Damaging | 0.50 |
| R20057C | Tolerable | Probably damaging | Disease causing | Medium | Damaging | Damaging | Damaging | 0.62 |
| P11728R | Tolerable | Benign | Polymorphism | Low | Tolerable | Tolerable | Tolerable | 0.22 |
| Y26509H | Tolerable | Probably damaging | Disease causing | Low | tolerable | Damaging | Tolerable | 0.50 |
| E30120K | Tolerable | Probably damaging | Disease causing | Neutral | Damaging | Damaging | Damaging | 0.68 |
| Q33877X | - | - | Disease causing | - | Damaging | - | Damaging | 1.0 |
| I19036T | Damaging | Possibly damaging | Disease causing | Medium | Damaging | Damaging | Tolerable | 0.45 |
| T25821M | Tolerable | Probably damaging | Polymorphism | low | Tolerable | Tolerable | Tolerable | 0.09 |

Abbreviations: AA, Amino Acid; SIFT, Sorting Intolerant From Tolerant;PolyPhen2, Polymorphism Phenotyping v2; FATHMM, Functional
Analysis through Hidden Markov Models; CADD, Combined Annotation Dependent Depletion;

### 3.4 Alignment of TTN protein sequence across species reveals highly conserved protein domains

We further narrowed down the list of 21 damaging variants to seven variants based on their rarity and downstream functions. A multi-species alignment of TTN protein in *H. sapiens, B. Taurus, M. musculus, R. norvegicus,* and *C. lupus* showed evolutionary conservation of residues D60, R22555, A14990, E24390, R18824, E30120 and Q33877 across species (see **Supplement Figure 1)**. Consurf prediction revealed that D60Y, A14990GfsTer4, R18842H, E24390KfsTer15, E30120K, and Q33877X were localized in the exposed region (**Table 3)**. Variations in the exposed regions can impact protein were more accessible for interactions with other molecules, which means they could significantly impact the protein’s role in cellular processes. On the contrary, R22555H residue was predicted as functionally significant indicating its involvement as an active site for downstream cellular pathways.

**Table 3:**
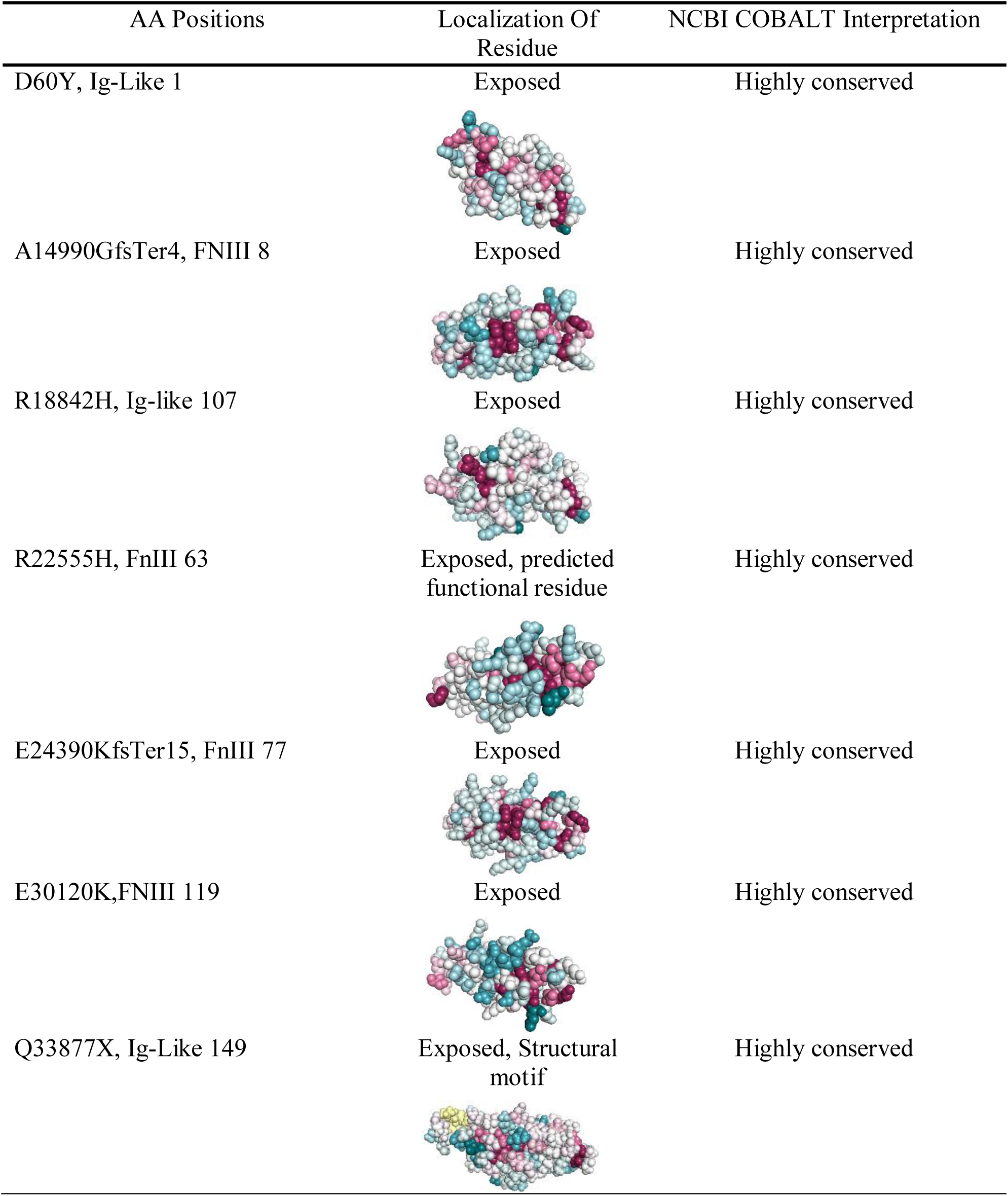
Phylogenetic conservation of TTN variants.

| AA Positions | Localization Of Residue | NCBI COBALT Interpretation |
| --- | --- | --- |
| D60Y, Ig-Like 1 | Exposed | Highly conserved |
| A14990GfsTer4, FNIII 8 | Exposed | Highly conserved |
| R18842H, Ig-like 107 | Exposed | Highly conserved |
| R22555H, FnIII 63 | Exposed, predicted functional residue | Highly conserved |
| E24390KfsTer15, FnIII 77 | Exposed | Highly conserved |
| E30120K,FNIII 119 | Exposed | Highly conserved |
| Q33877X, Ig-Like 149 | Exposed, Structural motif | Highly conserved |

### 3.5 Psipred predicts significant changes in the secondary structure of *TTN* variants

The PsiPred algorithm predicted that TTN would undergo significant changes in its secondary structure due to variants D60Y, R22555H, A14990GfsTer4, E24390KfsTer15, R18824H, E30120K and Q33877X. In the D60Y variant, the beta-strand became shorter upstream of the variation, potentially disrupting the protein’s structure and might affect its functional capabilities. Notably, in the A14990Gfs and E24390Kfs variants, the beta-strands present downstream of the variation site were completely abolished, suggesting aberrant protein structure possibly causing functionally abnormal protein-protein interactions. Additionally, in the nonsense variation Q33877X, the entire protein structure was distorted. (**Supplement figure 2**). Likewise, all the 21 abovementioned damaging variants demonstrated structural changes (data not shown).

### 3.6 Alpha-fold predicts protein models with a high confidence score

AlphaFold (AF) produces a per-residue estimate of its confidence on a scale from 0 - 100. This confidence measure is called pLDDT (predicted Local Distance Difference Test) and corresponds to the model’s predicted score on the lDDT-Cα metric. Based on the pLDDT scores, AF delivers 5 models on a ranking basis; higher the rank, higher the acceptance of the structure (**Figure 3**). The top-ranked model (Rank 1) based on AlphaFold pTM (predicted template modeling) score represents the highest accuracy level for each case. Therefore, Rank 1 structures were selected and equilibrated using MD simulations to further perform docking studies.

**Figure 3:**
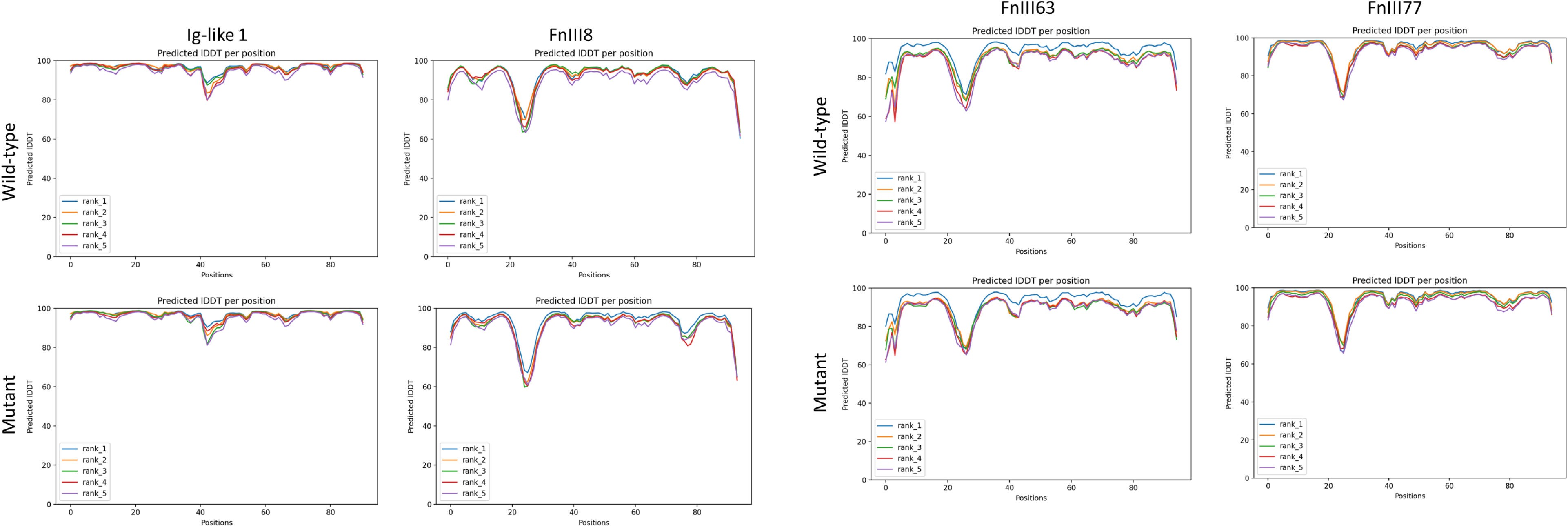
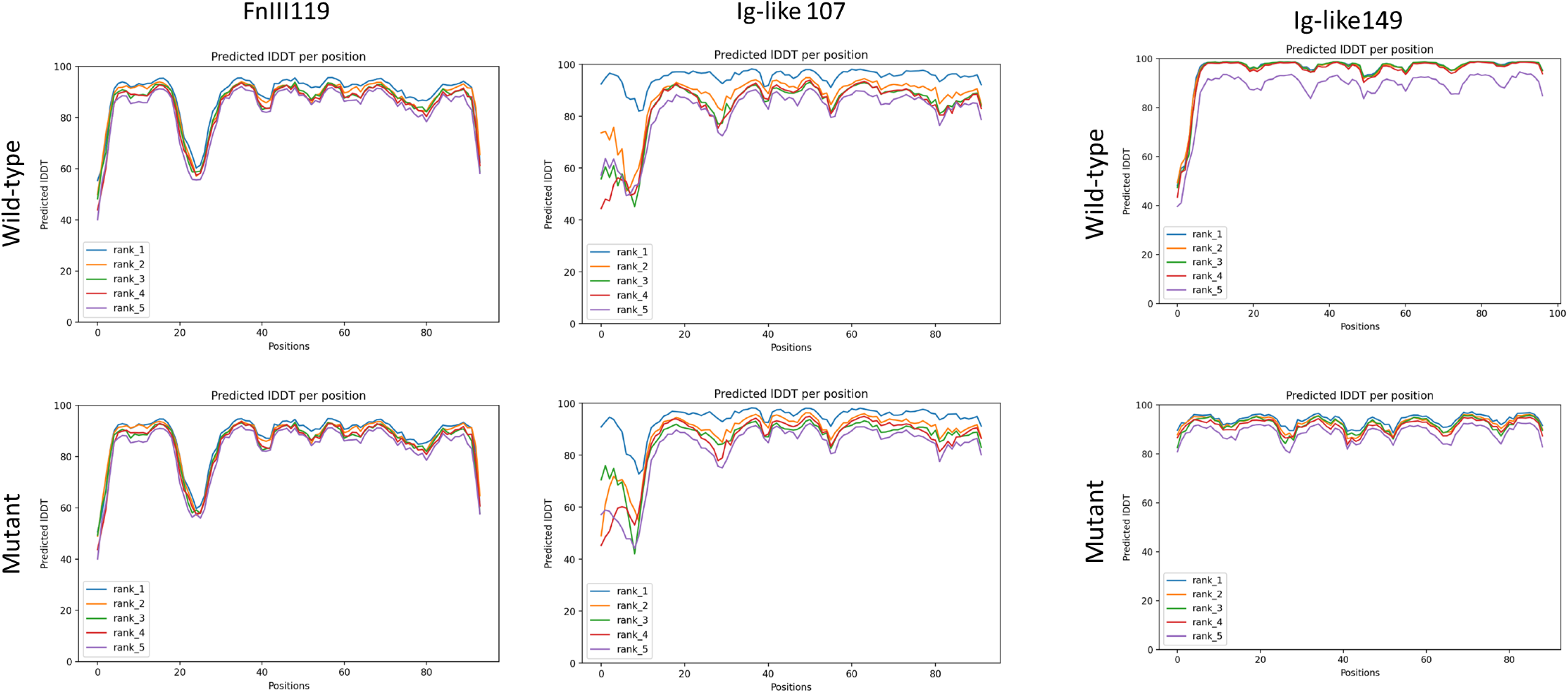
Representation of pLDDT plots of wild-type and mutant protein models obtained from Alpha-fold2googlecolab; AlphaFold2 automatically scanned the PDB for templates that were similar in sequence to the input FASTA sequence and produced a sequence coverage plot showing the number of homologs identified along the representative sequence and colored by the sequence identity of the homologs

### 3.7 Protein-protein docking reveals variations affect the molecular interactions of TTN with its interacting partners

We obtained 10-12 clusters, each containing four separate docked structures for every interacting partner. These structures are evaluated based on various parameters such as Z-score, cluster size, HADDOCK score, van der Waal energy, electrostatic energy and buried surface area (BSA). The clusters were ranked based on the average score of the top 4 members within each cluster and are numbered according to their size, indicating the number of members within each cluster. The differences in HADDOCK score (Supplement Table 3) were observed mainly due to restraint energy. We found R22555H, E24390Kfs, and Q33877X-WT TTN complexes showed more negative scores, indicating lower restraint energy compared to mutant complexes revealing their greater flexibility and tendency towards increased stability. The docking interactions between TTN domains and their interacting partners are provided in (**Figure 4).**

**Figure 4:**
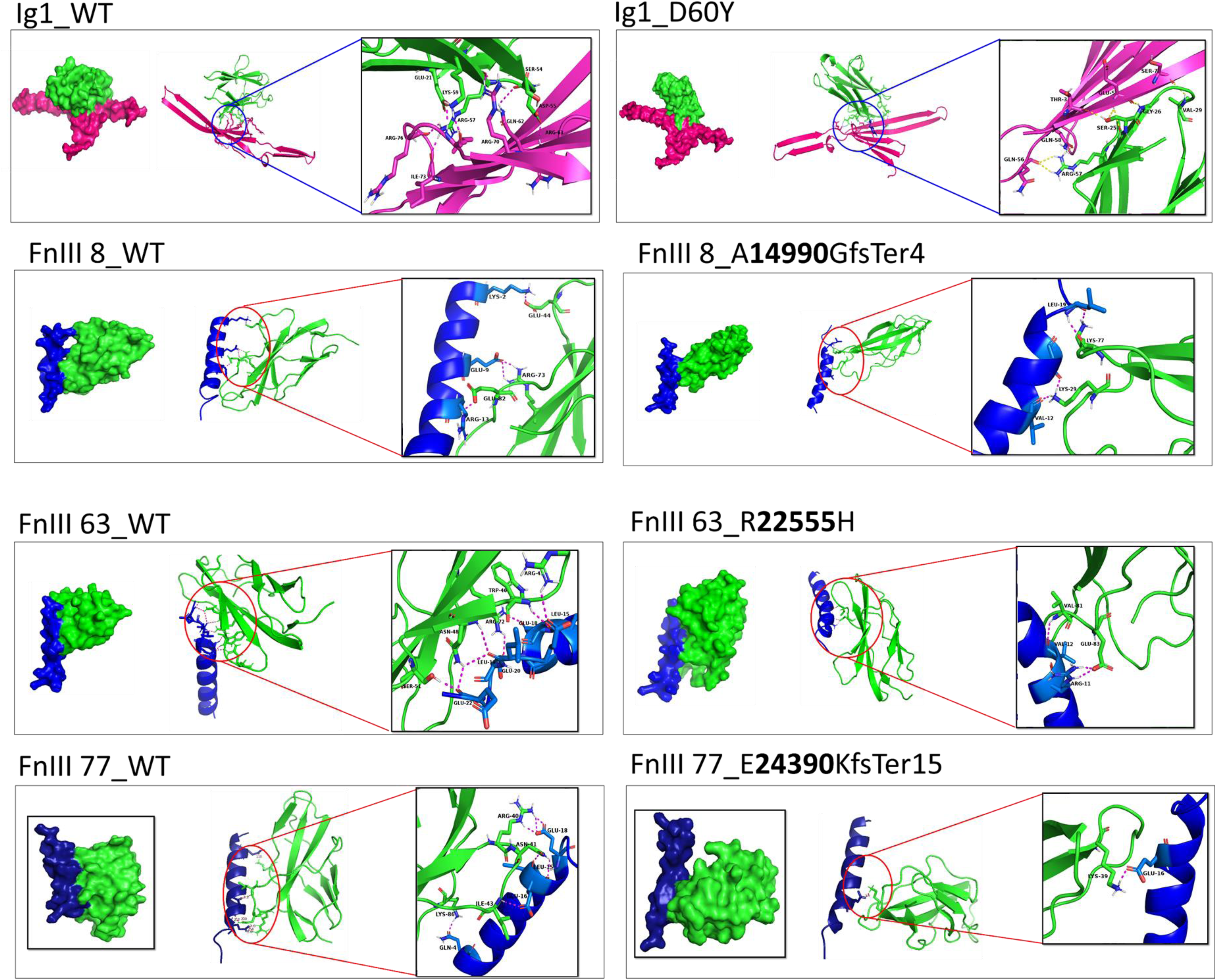
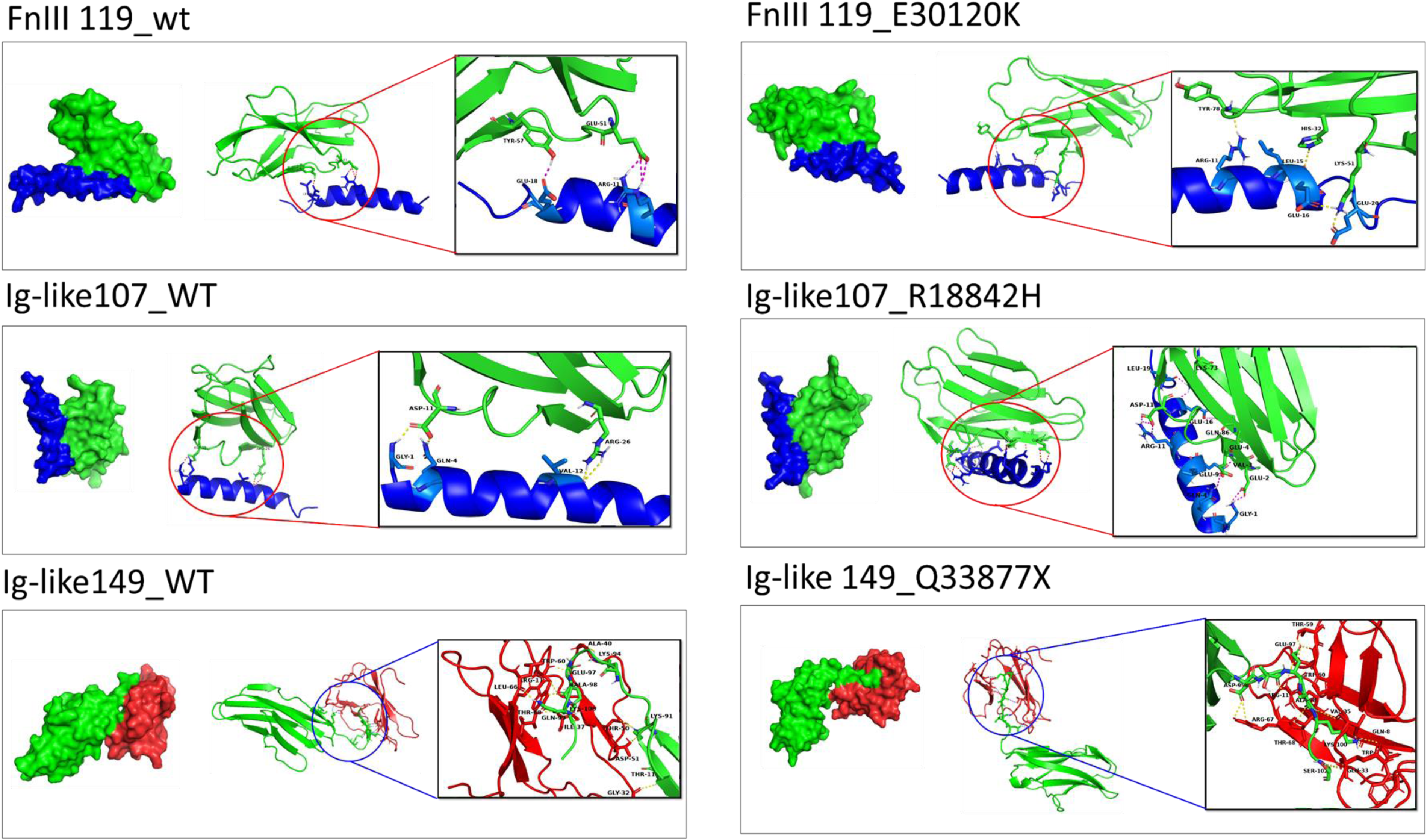
Protein-protein interaction of TTN protein with its interacting partners; We have shown protein-protein interactions of Wild-type TTN(Green) (Ig1-WT) as well as mutant TTN (D60Y) with TCAP (Magenta); FnIII 8, FnIII63, FnIII77, Ig-like107 and FnIII119 domains of wild-type as well as mutant TTN protein (Green) interacts with myosin (Blue); C-terminal Ig-like 149 domain of TTN (Green) interacts with Lamin (Red). The yellow/magenta colored interactions between residues represents hydrogen bonding. All the complexes depicted that both the interacting partners are forming numerous hydrogen bonds formed via various types of polar residues like ARG (Arginine), GLN (Glutamine), LYS (Lysine), ASP (Aspartic acid), SER (Serine), THR (Threonine), GLU (Glutamic acid) etc. which are showing significant role of hydrogen bonds in complex formation to increase binding affinity between them. The number of interactions between Fn63 and Fn77_WT-myosin was found to be more than their mutants which is in contrast to Ig-like 107_WT-myosin, which has more interactions in the case of mutant as compared to WT one. While there were comparable and similar numbers of hydrogen bonds formed with the rest of TTN domains with their interacting partners

### 3.8 Prediction of structural stability of protein complexes: findings from MD Simulation

In order to elucidate our findings, the results of the molecular docking study were evolved further using classical molecular dynamic simulation of 200 ns, to understand the structural stability of the formed complex system of various wild type and mutated TTN protein with TCAP, myosin and lamin and how its three-dimensional conformational changes evolve with respect to time.

#### 3.8.1 Comparison of RMSD, RMSF and RoG

The RMSD plot of the simulated system indicates that the Ig1-TCAP WT complex (purple) has an RMSD value of 0.6 to 2.5 nm for the first 25 ns after which eventually increased to 6 nm and was stable only for 25 ns. Similarly, for the Ig1-TCAP mutant complex (blue), the RMSD value was observed initially to be in the same range but eventually formed a more rigid structure than the WT one. The RMSD plot of another complex (Figure: 5) TTN FnIII8-Myosin WT (green) indicates an RMSD value of 0.25 to 0.75 nm during the initial 80 ns and rises up to 3.75 nm whereas for the FnIII8-Myosin mutant (blue) the RMSD value was 0.3 to 1.4 nm where the system seems to stabilize after the first 30 ns of the simulation and the RMSD remained constant at 1.1 nm which indicates that the system reached convergence and became more stiffer than mutant complex. Another myosin binding domain Ig-like107-Myosin complex indicates an RMSD value of 0.23 to 0.55 nm which seems to be stable throughout the trajectory as compared to mutant complex (blue) that showed 0.2 to 1.1 nm for the first 150 ns of the simulation after which the RMSD value increased up to 4.4 nm and was stable for 150 ns. FnIII 63_myosin WT complex (purple) have an RMSD value of 0.2 to 1.7 nm for the first 75 ns, after which it eventually increased to 7 nm. Similarly, for the FnIII 63_myosin mutant (blue) the RMSD value was 0.2 to 2 nm for the first 30 ns of the simulation after which the RMSD value increased up to 7.5 nm. The RMSD plot of another myosin-binding simulated system FnIII77-Myosin WT complex (green) have an RMSD value of 0.3 to 1.6 nm during the initial 100 ns of the simulation and then rises up to 4.5 nm as compared to mutant complex (blue) that showed a RMSD value of 0.2 to 0.7 nm for the first 75 ns of the simulation after which the RMSD value increased up to 2.7 nm. The RMSD plot of the FnIII 119-Myosin WT complex (purple) have an RMSD value of 0.2 to 0.5 nm which seems to be stable throughout the simulation. Similarly, for the Fn119-Myosin mutant (blue) the RMSD value was 0.2 to 1.5 nm for the first 90 ns of the simulation after which the RMSD value eventually increased to 6.5 nm. In the Lamin-binding domain, Ig-like 149-Lamin WT complex (green) have an RMSD value of 0.3 to 1.8 nm for the first 130 ns after which it increased up to 7.1 which was due to the breakdown of the complex. Similarly, for the Fn149_mutant (blue) the RMSD value was observed to be in the range of 0.2 to 1.4 nm for the first 50 ns of the simulation after which the RMSD value increased up to 5.1 nm and was stable only for 50 ns.

**Figure 5:**
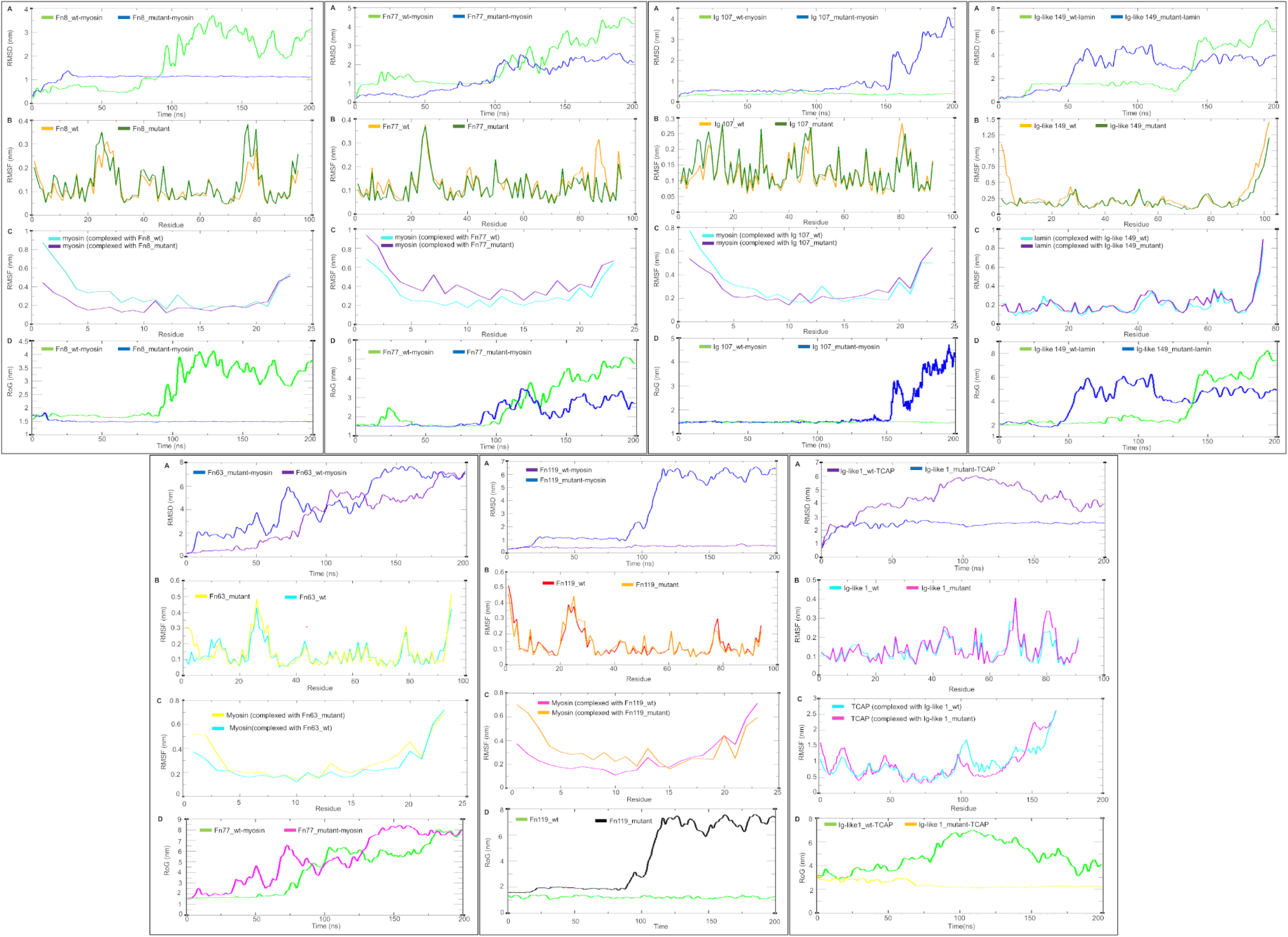
Molecular dynamics simulation analyses for the 200 ns simulation trajectory of the Titin (TTN) protein with its interacting partners;in each panel (A) represents the root mean squared deviation (RMSD) plots for the complexes TTN_WT with interacting partners (TCAP, Myosin and Lamin) and TTN_Mutant withInteracting partners; (B) shows root mean squared fluctuation (RMSF) (C) shows root mean squared fluctuation (RMSF) for the TTN_WT and TTN_Mutant complex; (D) illustrates the radius of gyration (RoG) plots for the complexes.

Throughout the MD simulation, the RMSF analysis was utilized to explain local flexibility changes at the atomic level. When compared to WT, all variations have large RMSF fluctuations at the atomic level. RMSF of Ig-like 1 WT, FNIII 8 WT, Ig-like 107 WT, FnIII 63 WT, and FnIII 119 WT complexes with their partners was greater than that of the mutant (Figure: 5). This suggested that these domains in the WT form exhibited more dynamicity and possibly more flexibility compared to their mutant counterparts. In contrast to the other domains, the FnIII 77 domain with myosin had a lower RMSF in the WT than in the mutant. This is an exception, suggesting that the variation in the FnIII 77-myosin complex increased its flexibility or fluctuation. The Ig-like149-lamin complex showed a lower RMSF in the WT compared to the mutant, similar to the FnIII 77-myosin complex. This indicates that the variation in the Ig-like149-lamin complex also leads to increased flexibility. In summary, most domains exhibited higher flexibility in their WT forms compared to their mutant counterparts, except for the FnIII 77 and Ig-like149 domains, where the mutants showed greater flexibility.

Another critical aspect, the RoG for monitoring the simulation trajectory, which helps to measure the distribution of the atoms of the protein around the three-dimensional space. The average RoG values (Figure:5) for the complex Ig1_WT-TCAP and the complex Fn8_WT-myosin (green) were in the range of 3-7nm and 1.5-4.2nm, respectively, which is quite larger than their respective mutant complex showing a stable RoG value of ∼2.7nm (yellow, Ig1_mutant-TCAP) and ∼1.52nm (blue, Fn8_mutant-myosin), indicating more tighter mutant complex. Conversely, the RoG value for the complex Ig-like 107_WT-myosin (green) was in the range of 1.45 to 1.58 nm having good stability and for Ig-like 107_mutant-myosin (blue) was in the range of 1.4-1.65 for the first 150 ns and then went up to 4.5 nm which was again due to the breaking of the complex formed. The RoG value for the complex Fn63_WT-myosin (green) was in the range of 1.6 to 2.7 nm for the first 75 ns, representing good stability after which it eventually increased to 8 nm due to the breakdown of the formed complex and for Fn63_mutant-myosin (magenta) was in the range of 1.6 to 2.5 nm for the first 30 ns and then went up to 8.5 nm which was again due to the breakdown of the complex formed. The RoG value for the complex Fn77_WT-myosin (green) was in the range of 1.6 to 2.5 nm for the first 100 ns and then went upwards to 5.2 nm, which is due to the myosin leaving the complex and for Fn77_mutant-myosin (blue) was in the range of 1.5-1.6 for the first 75 ns and then went up to 3.4 nm. Furthermore, the RoG value for the complex Fn119_WT-myosin (green) was in the range of 1.2 to 1.5 nm having good stability and for Fn119_mutant-myosin (black) was in the range of 1.6 to 2 nm for the first 90 ns and then went up to 8 nm which was again due to the breaking of the complex formed. The RoG value for the complex Ig-like 149_WT-lamin (green) was in the range of 2 to 2.8 nm for the first 130 ns having good stability after which it increased up to 8.4 nm and for Ig-like 149_mutant-lamin (blue) was in the range of 1.8 to 2.4 nm for the first 50 ns and then went up to 6.6 nm.

#### 3.8.2 Comparison of HBN and SASA analysis

The comparison of H-bond analysis between mutant and WT complexes revealed significant differences, highlighting the rigidity and compactness of the complexes. The D60Y mutant complex consistently maintained an average of 5 H-bonds while the WT complex had only 2 H-bonds (Figure 6). This indicates that the variation enhances stability by promoting stronger interactions. Similarly, the A14990Gfs, E30120K, and E24390KfsTer15 mutant complexes demonstrated a higher average number of H-bonds compared to its WT counterpart. Conversely, R22555H and R18842H mutants depicted lesser number of H-bonds as compared to their WT. While Ig like 149_WT and its mutant (Q33877X) showed same number of H-bonds with an average of 3. And all the H-bonds formed were observed to be lying within the cut-off distance 0.35nm (Figure 6) for all the complexes. Furthermore, to understand the hydrophobic core stability of the TTN protein structure, as it is a crucial parameter to be considered since the nonpolar amino acids corroborate the stability of the globular protein in solvent media by shielding the nonpolar amino acid residues inside the hydrophobic core of the protein so isolating them from the aqueous environment. We carried out SASA analysis on the trajectories (Figure 6), it was observed that the SASA of Ig1_wt-TCAP was 175 to 230 nm^2^ (Average 202 nm^2^) in comparison to the Ig1_mutant-TCAP which was in the range of 170 to 221 nm^2^ (Average 195 nm^2^). Likewise, Fn8_wt-myosin was a bit unstable in the range of 75 to 90 nm^2^ in comparison to the Fn8_mutant-myosin which was 75 to 85 nm^2^. In addition, the SASA analysis of the trajectories, suggested that Ig like 107_wt-myosin was in the range of 73 to 81 nm^2^ in comparison to the Ig like 107_mutant-myosin which was in the range of 78 to 92 nm^2^ which suggest it was more exposed to the solvent front. Likewise, the SASA analysis suggested that Fn63_wt-myosin was in the range of 84 to 93 nm^2^ in comparison to Fn63_mutant-myosin which was in the range of 77 to 98 nm^2^. Moreover, Fn77_wt-myosin was in the range of 77 to 93 nm^2^ whereas for Fn77_mutant-myosin SASA was in the range of 76 to 96 nm^2^ and was seen to be more fluctuating as compared to the Fn77_wt-myosin complex. Likewise, the SASA of Fn119_wt-myosin was 71 to 90 nm^2^ in comparison to the Fn119_mutant-myosin which was in the range of 75 to 94 nm^2^. Likewise, the SASA analysis of the trajectories, suggested that Ig like 149_wt-lamin was 107 to 129 nm^2^ in comparison to the Ig like 149_mutant-lamin which was in the range of 105 to 127 nm^2^.

**Figure 6:**
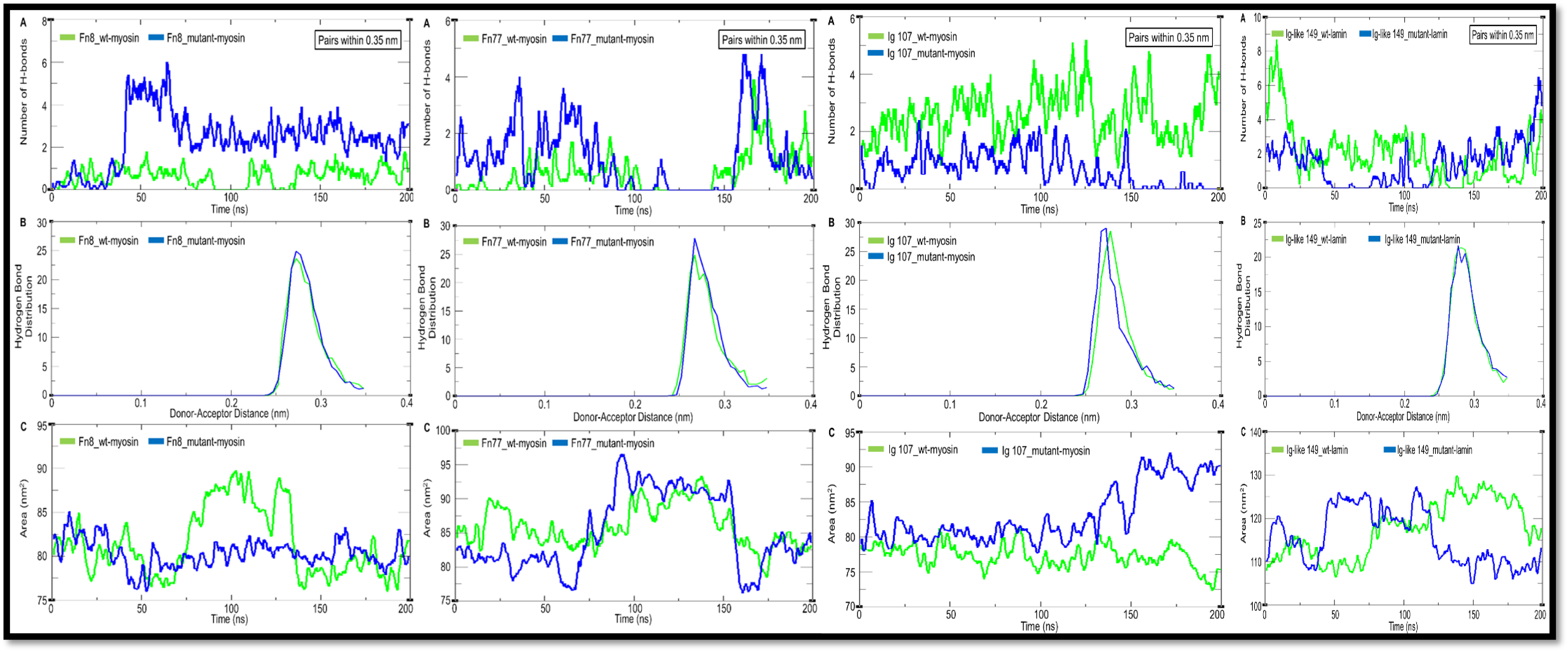
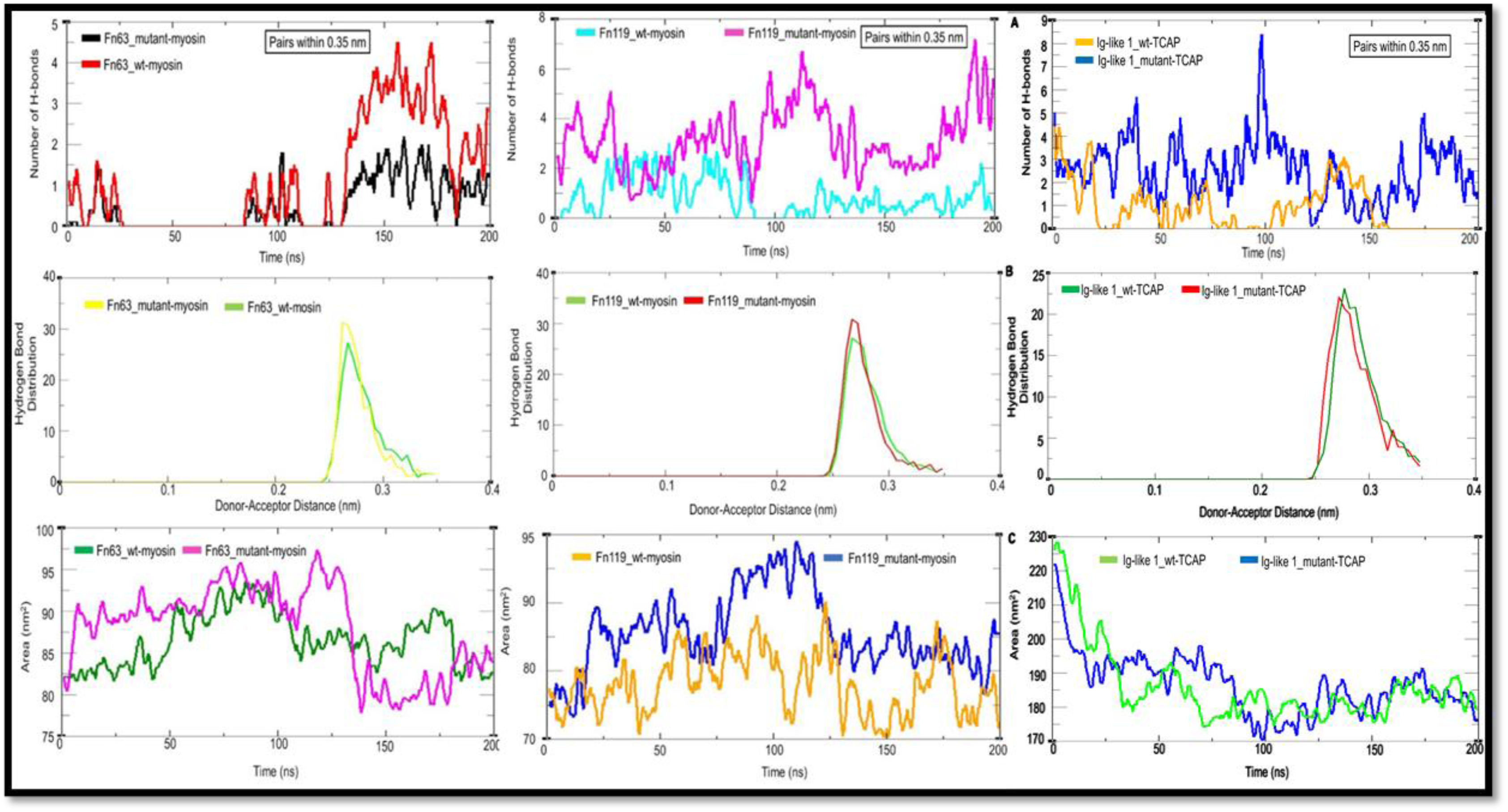
Hydrogen bond landscape and solvent accessible surface area analysis of the Titin (TTN) protein, WT and Mutant complexes. (A) shows the average number of hydrogen bonds formed between the TTN-WT complex and TTN-Mutant complex; (B) illustrates the average hydrogen-bond distance maintained which was found to be around the cut-off for donor–acceptor distance that was set at 0.35 nm. (C) denotes the change in total solvent accessible surface area (SASA) of the system with respect to time under simulation.

The eigenvalues of the first 10 principal components (eigenvectors) were plotted for WT as well as mutant complexes. It was observed from the graphs that The first two principal components were crucial in explaining structural changes, indicating different geometric conformations leading to energy minima states. Overall, mutants exhibited diverse characteristics compared to WT complexes, influencing their stability and conformation (Figure 7).

**Figure 7:**
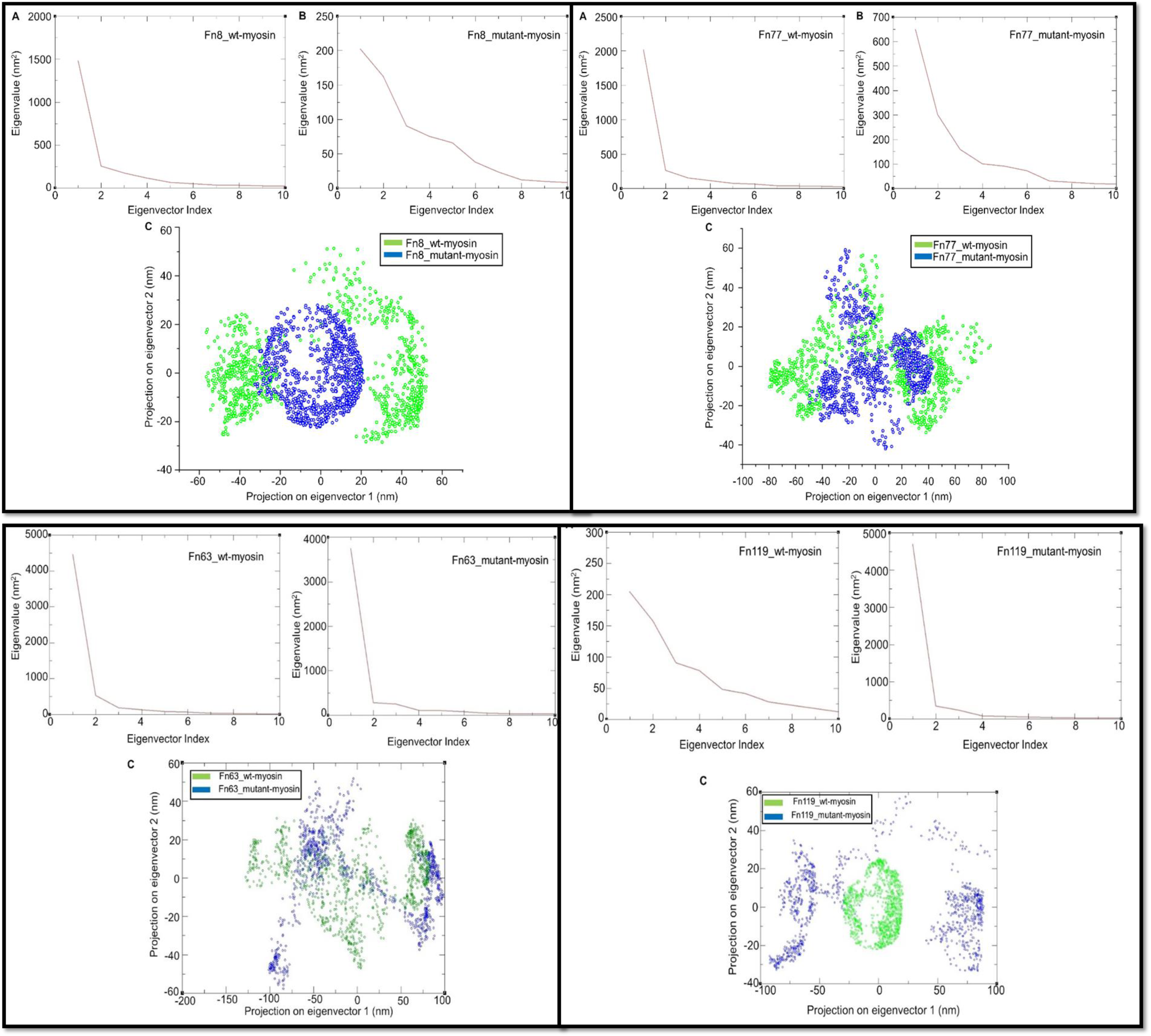
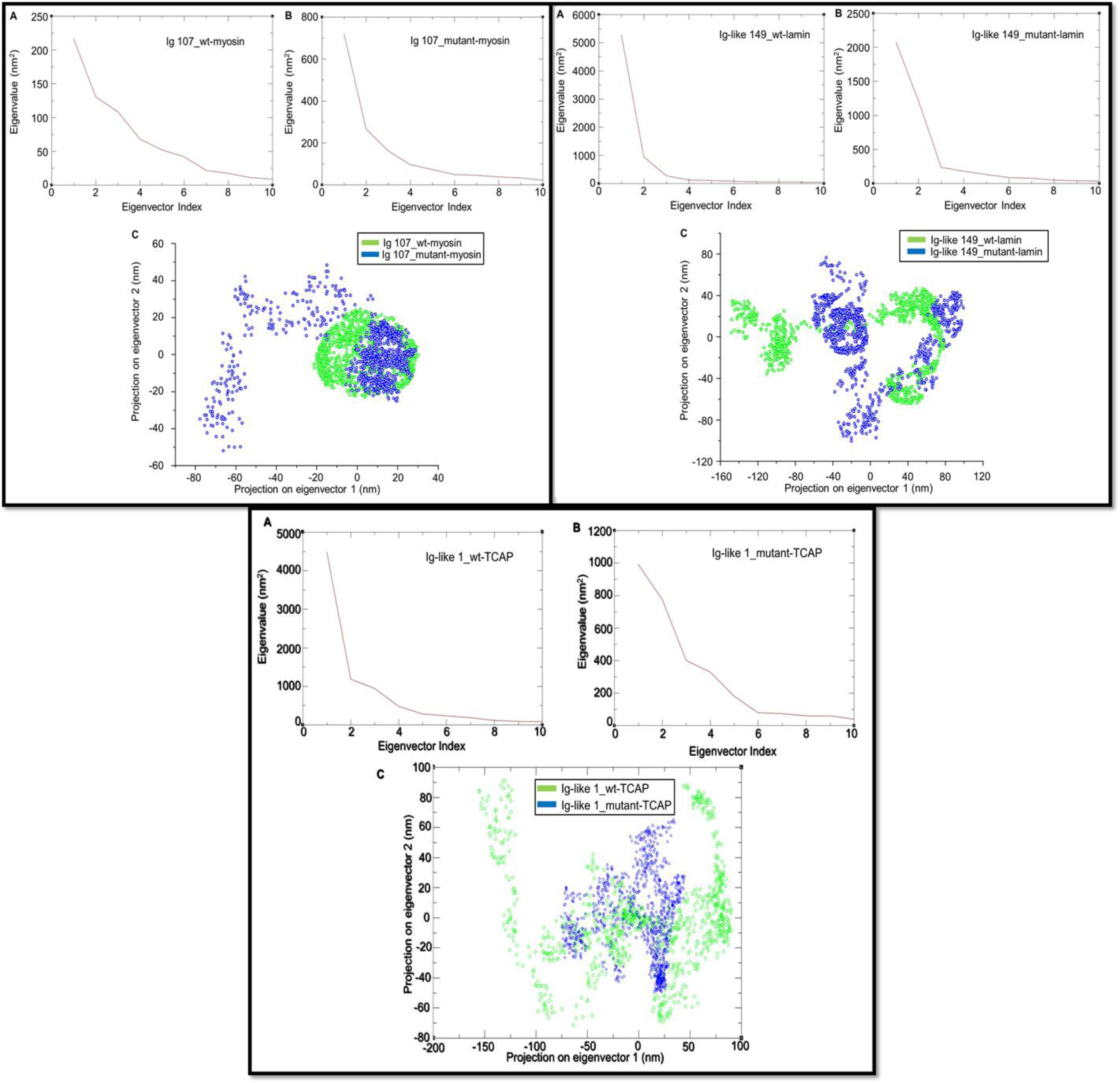
(A) & (B) in all the plots represent the eigenvalues of the first 10 principal components for the TTN-WT complex and TTN-mutant complex respectively. The analysis suggests that the first two principal components eigenvector accounts for most of the motions of the system, while from the third component of eigenvector the contribution seems insignificant.; (C) The first two PCs were used to generate the 2D projections of TTN_WT complex (green) and TTN_mutant complex (blue) by plotting it on x and y axis in order to explain the data points. Where, each dots represents a specific conformational state of the system that it underwent during the 200 ns simulation.

#### 3.8.3 Free-Energy Landscape Analysis

To visualize the energy minima landscape of the complex systems, we used the RMSD and RoG as the two reaction coordinates, revealing the changes in its Gibbs free energy (ΔG) values ranging between 0 and 14.70 kJ/mol (Figure 8) indicated by the colour coding of purple as the lowest energy minima and red being the highest energy respectively. The shape and size of the funnel formed in the 3D plot indicates the stability of the system. Smaller and more centralized blue areas in the 2D plot represent the complex within the cluster with the highest stability. It was observed that Fn8_wt-myosin and Ig like 1_wt-TCAP complex formation was not much stable as compared to the Fn8_mutant-myosin and Ig like 1_mutant-TCAP complex which has a single funnel shape in the 3D FEL plot denoting a stable folding process. Likewise, the Fn77_wt-myosin and Fn77_mutant_myosin seem to have some uncertainties where the funnel distribution is more widespread in the Fn77_mutant-myosin as observed from the 2D FEL plot indicating comparatively more stability of the Fn77_wt-myosin complex. Similarly, it can be seen that Ig like 107_wt-myosin, Fn63 wt-myosin and Fn119_wt-myosin has a more stable folding process as compared to the Ig like107, Fn63 and Fn119 _mutant-myosin complex. Also, the Ig like 149_wt-lamin complex was having better folding process to reach energy minima as compared to the Ig like149_mutant-lamin.

**Figure 8:**
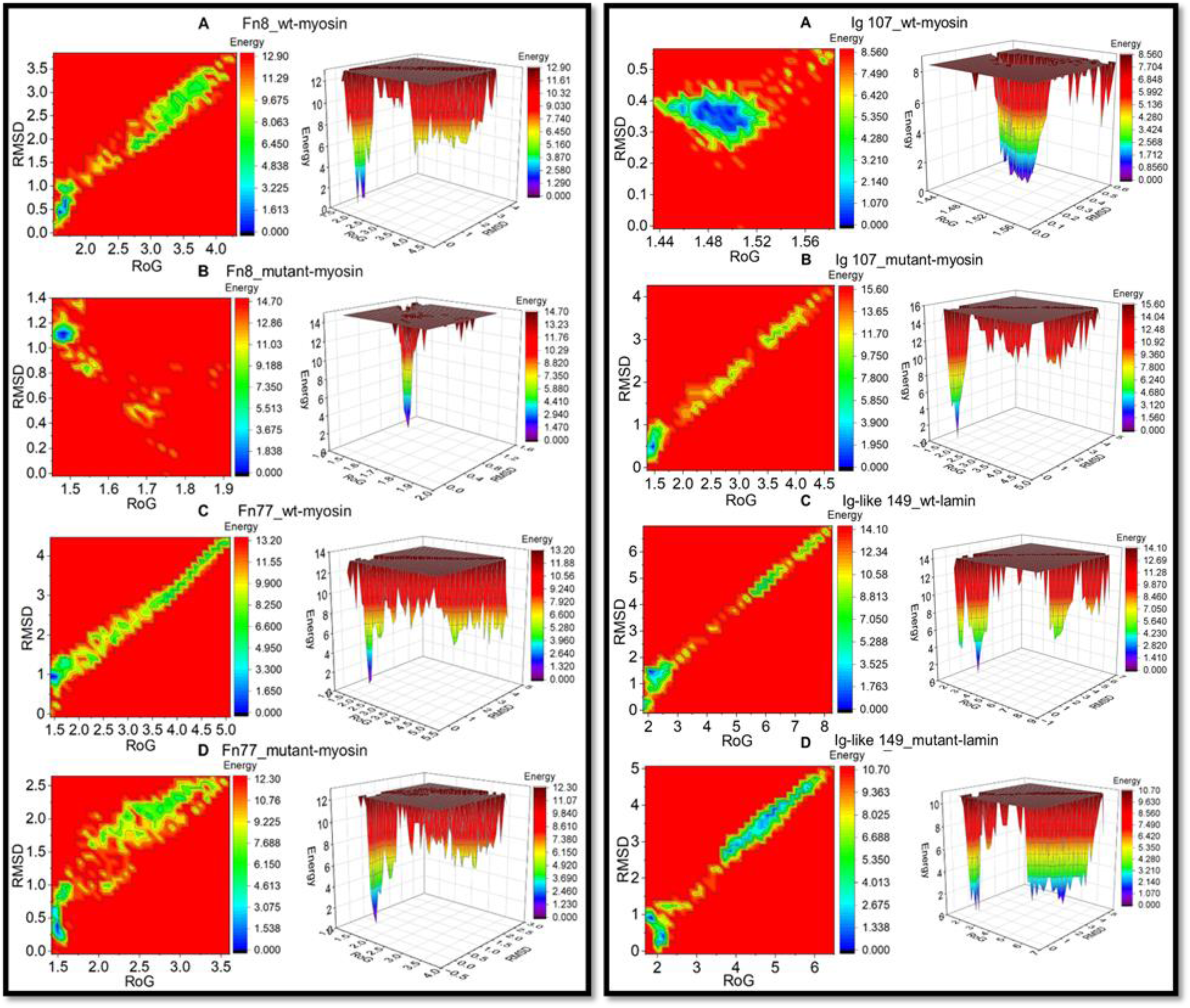
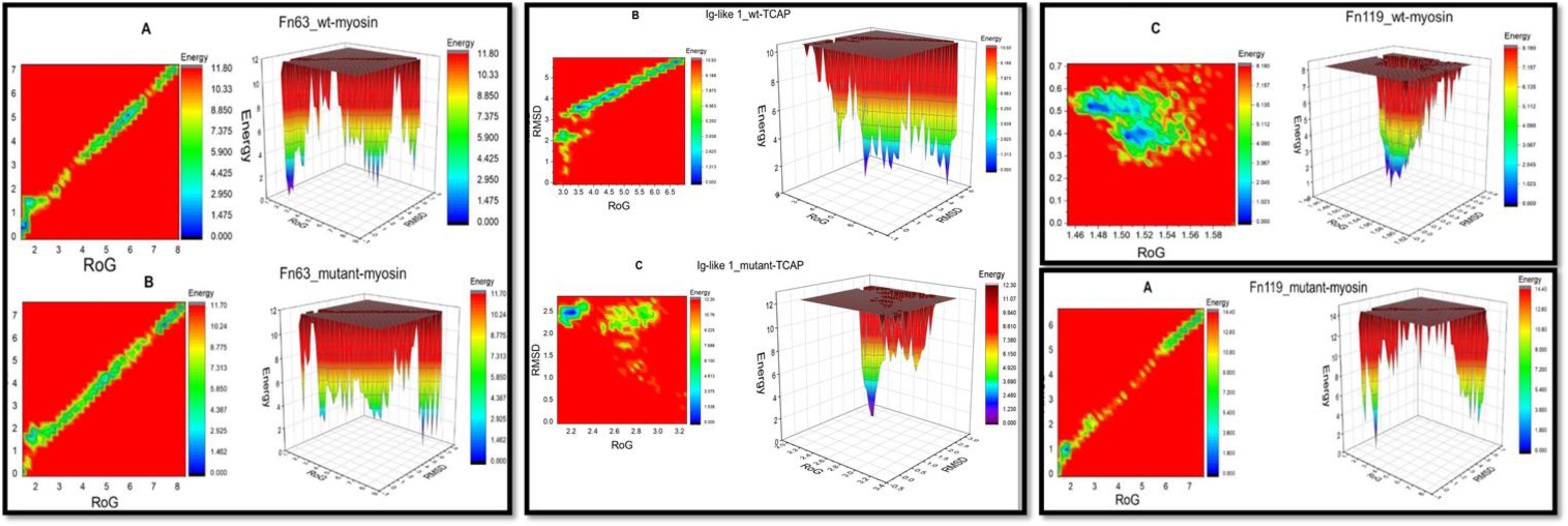
The 2D and 3D free energy landscape diagrams as a function of RMSD and RoG as the two reaction coordinates that depict the folding of the Titin (TTN) protein. The free energy is displayed in terms of kJ/mol in the indicated color codes where the purple color denotes the lowest energy and the red color denotes the highest energy. The simulated folding of the protein in the process of forming a stable system forms a distinct narrow funnel shown in 3D diagram that ultimately denotes a stable folding state with lowest energy minima and thereby highest stability. The 2D contour plots and the 3D projections were plotted using OriginPro 2017 software (OriginLab Inc., Northampton, MA, USA).

### 3.9 Protein stability prediction

Our analysis revealed that three variants, namely D60Y, A14990Gfs, and R22555H, exhibited the highest degree of instability across all protein stability prediction tools utilized (Table 5). Based on this screening, the variants were categorized into two groups based on their impact on protein stability: those that were destabilizing (including R18842H, E24390Kfs, E30120K, and Q33877X) and those that were mostly destabilizing (comprising D60Y, A14990Gfs, and R22555H). Analysis of physicochemical parameters revealed theoretical isoelectric point (pI), hydrophobicity, and polarity of protein complexes fluctuate with the variants. Increased hydrophobicity was shown by all myosin-binding TTN-MYH7 complexes, whereas, TCAP and LMN binding complexes shown more hydrophilic nature (Detail comparison is shown in Table 6).

**Table 4:**
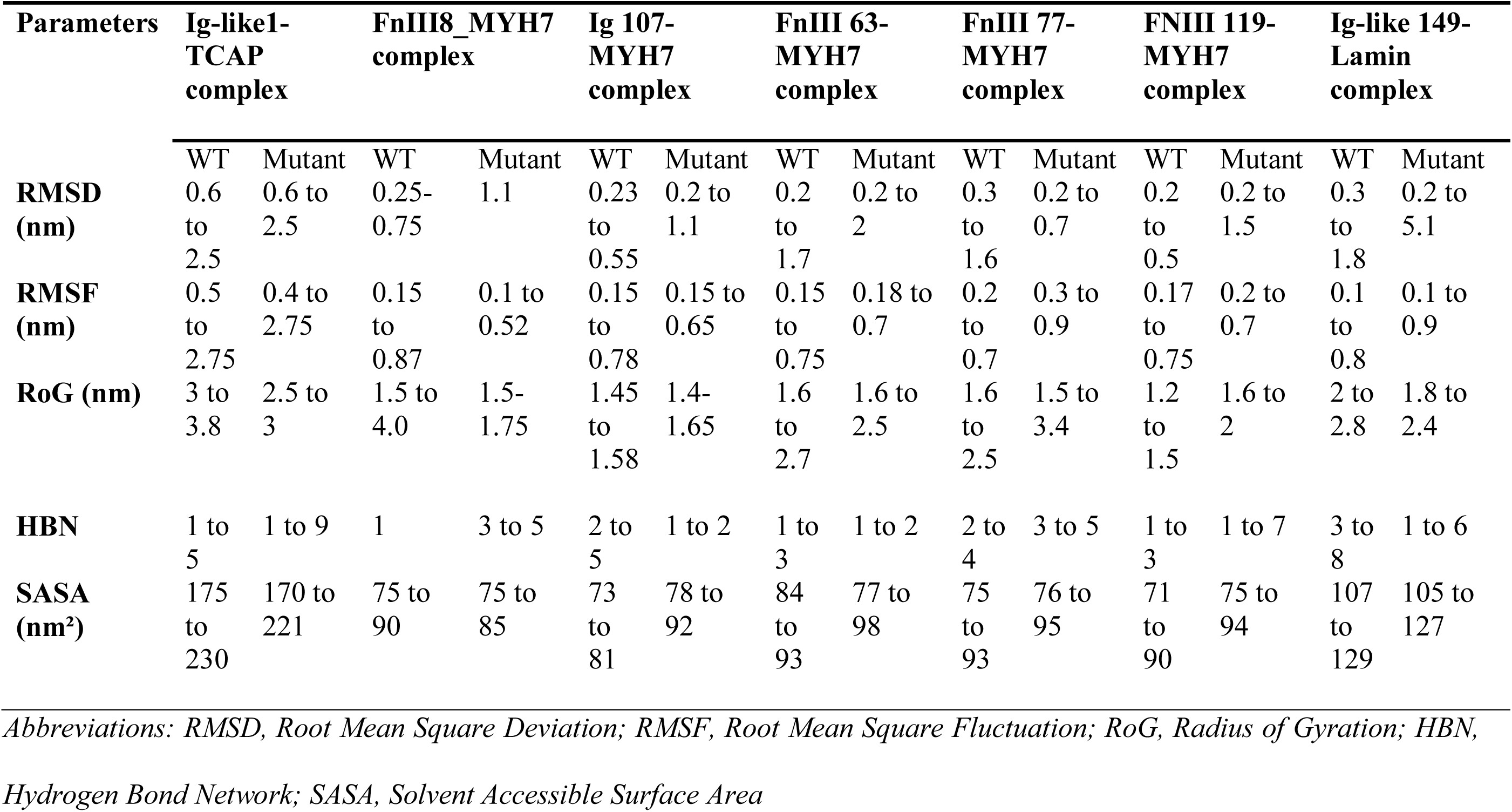
Interpretation of MD simulation result (RMSD, RMSF, RoG, HBN, SASA analysis)

**Table 5:** Prediction of protein stability of TTN damaging variants.

| AA change | DYNAMUT2 | MUpro | CUPSAT | DUET | I-mutant | SDM | mCSM |
| --- | --- | --- | --- | --- | --- | --- | --- |
| D60Y, (Ig-Like1) | Destabilizing, -0.49kcal/mol | Destabilizi<br>ng, -<br>0.85<br>kcal/mol | Destabilizin<br>g, -<br>1.33<br>kcal/mol | Destabilizin<br>g,<br>-0.831<br>kcal/mol | Destabilizin<br>g,<br>-1.13<br>kcal/mol | Destabilizin<br>g, -<br>0.67<br>kcal/mol | Destabilizin<br>g, -<br>0.719<br>kcal/mol |
| A14990GfsTer4, (FnIII 8) | Destabilizing, -0.56kcal/mol | Destabilizi<br>ng, -<br>1.36<br>kcal/mol | Destabilizin<br>g, -0.7<br>kcal/mol | Destabilizin<br>g, -<br>0.59<br>kcal/mol | Destabilizin<br>g,<br>-1.56<br>kcal/mol | Destabilizin<br>g, -<br>0.64<br>kcal/mol | Destabilizin<br>g, -<br>0.735<br>kcal/mol |
| R18842H, (Ig-Like 107) | Destabilizing, -1.19kcal/mol | Destabilizi<br>ng, -<br>1.28<br>kcal/mol | Stabilizing,<br>0.66<br>kcal/mol | Destabilizin<br>g, -<br>1.516<br>kcal/mol | Destabilizin<br>g,<br>-1.50<br>kcal/mol | Stabilizing,<br>-<br>0.35kcal/mo<br>l | Destabilizin<br>g, -<br>1.74<br>kcal/mol |
| R22555H, (FnIII 63) | Destabilizing, -1.26 kcal/mol | Destabilizi<br>ng, -<br>1.00<br>kcal/mol | Destabilizin<br>g, -4.44<br>kcal/mol | Destabilizin<br>g, -<br>2.176<br>kcal/mol | Destabilizin<br>g,<br>-2.08<br>kcal/mol | Destabilizin<br>g, -<br>0.82<br>kcal/mol | Destabilizin<br>g, -<br>1.992<br>kcal/mol |
| E24390KfsTer15, (FnIII 77) | Destabilizing, -1.26 kcal/mol | Destabilizi<br>ng, -<br>1.20<br>kcal/mol | Stabilizing,<br>2.1 kcal/mol | Destabilizin<br>g, -<br>1.199<br>kcal/mol | Destabilizin<br>g,<br>-<br>0.83kcal/mo<br>l | Destabilizin<br>g, -<br>0.15<br>kcal/mol | Destabilizin<br>g, -<br>1.42<br>kcal/mol |
| E30120K, (FnIII 119) | Stabilizing, 0.18kcal/mol | Destabilizi<br>ng, -<br>0.91<br>kcal/mol | Destabilizin<br>g, -0.53<br>kcal/mol | Destabilizin<br>g, -<br>0.285<br>kcal/mol | Destabilizin<br>g,<br>-0.74<br>kcal/mol | Destabilizin<br>g, -<br>0.18<br>kcal/mol | Destabilizin<br>g, -<br>0.56<br>kcal/mol |
| Q33877X, (Ig-Like 149) | -- | -- | -- | -- | -- | -- | -- |

Abbreviations: Dynamut2, Dynamic Mutant 2; MUpro , Mutant Protein Predictor; CUPSAT, Cologne University Protein Stability Analysis Tool; DUET, Double Mutation Estimator Tool; I-mutant, Internet-based Mutant; SDM, Site-Directed Mutator; mCSM, mutation Cutoff Scanning Matrix.

**Table 6:** Analysis of Physicochemical Parameters of TTN damaging variants.

| Parameters | Ig-like1-TCAP complex |  | FnIII8_MYH7 complex |  | Ig 107-MYH7 complex |  | FnIII 63-MYH7 complex |  | FnIII 77- MYH7 complex |  | FnIII 119-MYH7 complex |  | Ig-like 149-Lamin complex |  |
| --- | --- | --- | --- | --- | --- | --- | --- | --- | --- | --- | --- | --- | --- | --- |
|  | WT | Mutant | WT | Mutant | WT | Mutant | WT | Mutant | WT | Mutant | WT | Mutant | WT | Mutant |
| Theoretical pI | 7.18 | 8.77 | 5.70 | 6.15 | 9.33 | 8.98 | 5.83 | 5.64 | 6.36 | 10.00 | 8.24 | 8.96 | 8.96 | 8.69 |
| Polarity | Neutral | Decrease polarity | Acidic | Decrease polarity | Basic | Increase polarity | Acidic | Increase polarity | Acidic | Decrease polarity | Basic | Decrease polarity | Basic | Increase polarity |
| GRAVY | 0.004 | 0.029 | -0.722 | -0.683 | -0.341 | -0.327 | -0.178 | -0.164 | -0.562 | -0.465 | -0.306 | -0.311 | -0.334 | -0.307 |
| Hydrophobicity | - | Decreases | - | Increases | - | Increases | - | Increases | - | Increases | - | Decreases | - | Increases |
| Positive residue | 6 | 6 | 13 | 11 | 16 | 15 | 11 | 10 | 11 | 7 | 12 | 13 | 14 | 13 |
| Negative residue | 6 | 5 | 15 | 12 | 13 | 13 | 12 | 12 | 12 | 3 | 11 | 10 | 11 | 11 |

Abbreviations: pI, Isoelectric point; WT, Wild type; GRAVY, Grand Average of Hydropathy;

### 3.10 Structural evaluations

The Ramachandran plot analysis conducted using PROCHECK indicates that over 90% of the residues in all the modeled structures fell within the favoured regions. This high percentage of residues in the preferred regions suggested that the structures were of high quality and could be considered reliable models. This level of compliance with the favoured regions in the Ramachandran plot reflected proper folding and minimal steric clashes, supporting the overall validity of the modeled structures (Supplementary Figure 4). Additionally, PDBsum provided detailed information on the specific residues involved in interactions between the protein chains. We predicted interacting residues in WT-TTN domains (Supplement Figure 5).

## 4. Discussion

In the present study, we used WES technology to identify potential disease-causing variants in *TTN*. We have identified a total of 88 coding variants, of which 21 were pronounced as ‘damaging’. Among these 21 variants, 7 were prioritized for further downstream characterization.

In this investigation, all 15 probands harbour ample heterozygous *TTN* variations. The majority of them except one patient (SDCM1), carry multiple variations. Although 67 out of total 88 variations are assessed as benign, 21 variations (showing high damaging score) marked as deleterious, are proportionately associated with both FDCM and SDCM cases. Analysis of family members in FDCM cases has revealed majority of variations are inherited from any one of the unaffected parents, thus indicating intriguing nature of these variations. Interestingly, patients harbouring more than one damaging variation also show more severe phenotype, indicating possible effect of compound heterozygosity. In a comparable study, hypermutability of TTN gene in DCM cohort has been reported [37] where they have identified 348 missense variants in 37 probands out of 147 DCM cases analysed. Of these, 44 variants were classified as severe variants. Similarly, in another European cohort study including 639 DCM patients [38] *TTN* variations are identified in 19% familial and 11% sporadic cases, depicting high incidence of TTN variations in DCM patients. To date, several truncation variations in the *TTN* gene have been predominantly linked to DCM [8, 39]. However, detection of *TTN* truncating variants in healthy individuals in the general population raises questions about the clinical significance of these variations [40]. Therefore, it is crucial to estimate the frequency of the variants in the general population and in controls to comprehend the clinical implications of specific variants. We discern that the MAF of all 21 ‘damaging’ variants is extremely low (<0.05) in ExAc and gnomAD databases. However, two variants namely S23807N (MAF=0.079273) and A15520V (MAF=0.072684) show moderately higher MAF in 1000Genomes (Supplementary Table 2), where all other variants have MAF<0.05. Further, in the Indian population databases (IndiGenomes, Index-db, Genome Asia), the MAF of all the 21 variants is exceptionally low (<0.1%). This underscores the rarity of these variants in healthy individuals and supports the notion of their potential pathogenic impact.

Out of 21 deleterious variants, majority are localised to A-band region of TTN, along the line of total variations recorded, i.e., 44.3 % in the A-band region, compared to 37.5% in the I-band, 10.2% in the M-band, and 7.9% in the Z-disc region. Comparable higher clustering of variants in A-band region has been observed in the previous investigation [37]. Similarly ‘overrepresentation’ of truncating variations in A-band region, which are absent from Z-disk and M-band domain has been reported by Herman et al., (2012) [8]. This study is also supported by the observation of Pugh et al., (2014) [41], where they have also recorded *TTN* truncating variations are more prevalent in the A-band region compared to the Z-disk and M-band sections in DCM patients. Although no strong association of these variations with the risk of DCM could be established, another study argue that conserved regions in the A-band domain are critical for muscle function and variations can lead to dilated cardiomyopathy and other muscle diseases [39]. In our study, the variants are disproportionately clustered within the A-band region of TTN, specifically concentrated in the super-repeats region of the A-band that interacts with myosin. Incidentally, 3 out of 4 novel variants, namely, R18842H, A14990GfsTer4 and E24390KfsTer15 are located in this region. In previous studies, the FnIII domains have already been demonstrated to engage with myosin subfragment-1, which play a crucial role in the force generation on muscle length [42]. Therefore, we posit that missense and frameshift/ truncation variations within A-band could potentially exert functional detrimental effect on its binding partner myosin. Notably, our investigation has revealed that 60% of variants are concentrated within the Ig-like domain and 32% of variations are situated in the FnIII domain of TTN. This comprehensive distribution of subdomains in full-length TTN emphasizes its complex and modular nature. Such complexity is essential for the multifaceted functions of TTN in muscle physiology, viz., providing structural support, imparting elasticity, and regulating muscle contraction.

Analysis of the evolutionary conservation of amino acids is valuable for pinpointing variations associated with diseases, as disruptions in conserved amino acids are more prone to cause pathological conditions [43]. Further, protein-protein interfaces also reveal that conserved residues are often located at the core of interaction sites, ensuring the stability and specificity of these interactions [44]. In this study all 21 variants are predicted to be damaging, are localized in highly conserved regions of TTN, indicating their disease-causing possibility. Amino acids demonstrating a high degree of conservation across species are more likely to play a crucial role in the protein’s structure or function. Modifications to these conserved residues may result in significant functional consequences. For example, in the D60Y variant, the beta-strand preceding the variation site became shorter, which could potentially disrupt the protein’s functional capabilities. Since, beta-strands are considered critical components of the protein’s secondary structure, any alterations in their length or integrity can significantly affect the overall stability and function of the protein. The shortening of the beta-strand upstream of the D60Y variant may lead to improper folding or loss of structural rigidity, compromising the protein’s ability to perform its normal functions. Similarly, in case of A14990Gfs and E24390Kfs variants, the complete abolition of the beta-strands, located downstream of the variation sites, likely leads to a significant disruption in the protein’s architecture, rendering its inability to interact properly with other proteins. For the nonsense variation Q33877X, introduction of a premature stop codon caused a truncation of the protein, resulting in a severely distorted overall structure.

Moreover, the physicochemical characteristics of proteins, including their charge, hydrophobicity, and structure has been shown to be frequently undergoing substantial changes as a consequence of disease-causing variants [45]. Therefore, here we examine how variations impact protein structure, focussing on the physicochemical parameters, as these are the primary determinants of protein structure and function. Hydrophilicity is shown to be greater in mutant complexes of A14990Gfs, E24390Kfs, R18842H, E30120K and Q33877X. However, D60Y is showing lesser hydrophilicity among all, indicating this mutant complex would have more hydrophobic regions, which could affect its folding, stability, and interactions with other molecules (Table 6). In another study, Tokmakov et al., (2021) discussed the importance of ionizable groups and overall net charge of a molecule that can be significantly influenced by its three-dimensional structure and the pH of the surrounding environment [46]. We have also noted that the R22555H and A14990Gfs variations result in a more acidic nature, whereas the E24390Kfs variant exhibit extreme basic properties compared to the WT. Understanding of these changes are crucial for deciphering the functional implications of variations in proteins. Modifications in the physiological parameters of a protein due to variations can have profound effects on its structure, function, and role within the cell. These changes can lead to a range of biological consequences, from altered metabolic pathways to the development of diseases.

Our study is further focused on the *in-silico* molecular docking of TTN, particularly with its interacting partners to determine the impact of variations on inter-atomic interactions. We observed that the wild-type (WT) TTN complexes (either with Myosin or Lamin) formed more stable interactions, compared to the mutants, namely R22555H, E24390Kfs and Q33877X complexes (Figure 4). Visualization of the protein-protein complexes reveal greater interatomic synergy in the aforementioned WT complexes than in the mutant ones. The robustness of the docking results is thoroughly examined through extensive molecular dynamics simulation lasting 200ns. In a similar study, Lee et al., (2009) investigated the interactions between TTN and other sarcomeric proteins, such as actin and myosin. MD simulations have highlighted the critical interfaces and binding affinities, elucidating how TTN contributes to the overall stability and function of the sarcomere [47]. In another study, Suganthi et al., (2024) conducted a protein-protein interaction study between GAPDH and Chitinase using the HADDOCK tool. Their findings reveal that a more negative HADDOCK score indicates a more favourable docking interaction, suggesting a more stable and potentially biologically relevant complex [48]. MD simulations aim to mimic the authentic behaviour of molecules within their respective environments, considering their flexibility, dynamic movements, and evolving motions over time.

Our analysis parameters include RMSD, RMSF, RoG, H-bond number (HBN) and assessment of SASA. RMSD measures the average deviation of a protein’s atomic positions from a reference structure over time [49]. As shown in Figure 6, the myosin binding Ig-like 107 and FnIII 119-WT structure have showed the most stable complex as compared to their mutants. Among the mutant complexes, the A14990Gfs, and D60Y mutant remain structurally more stable. The lamin-binding Q33877X and myosin-binding R22555H mutants are stabilized only at the earlier time period of MD. Another myosin-binding E30120K mutant indicate extremely unstable complex throughout simulation. Thus, the RMSD analysis of the wild type and the mutants is resulted in a decrease in stability of TTN-mutant complex, except D60Y, E24390Kfs and A14990Gfs mutant complexes.

Theoretically, increased RMSF implies decreased stability of proteins [50]. Mutant D60Y and A14990Gfs proteins exhibit increased rigidity compared to their WT complexes. This observation is aligned with the RMSD values, emphasizing the structural stability of TTN domains imparted by these variations. Conversely, the variations (R18842H, E24390Kfs, R22555H, and E30120K) in myosin-binding domains, along with the variation in lamin-binding domain (Q33877X), demonstrate greater flexibility in mutant complexes than WT. This heightened flexibility is intricately linked to their interactions with other proteins, facilitating dynamic binding and functional versatility. The observed increase in rigidity in the mutant complex of D60Y and A14990Gfs further lead to reduction in the protein’s overall dimensions studied by RoG. RoG values for both of these mutant structures is found to be lesser than the WT-complex suggesting the structural changes induced by the variation cause the protein to adopt a more compact conformation. This compactness could influence the protein’s functional dynamics, potentially altering its interaction capabilities with other molecules. In other variants (R18842H, E24390Kfs, R22555H, E30120K, and Q33877X), the RoG values are comparable to or even exceeded those of the WT complexes, indicating a similar level of structural compactness. This similarity in RoG values suggests that these variants (R18842H, E24390Kfs, R22555H, E30120K, and Q33877X) maintain a compact structural configuration akin to that of the WT. Such findings imply that the overall folding and spatial arrangement of these protein variants are not significantly disrupted, allowing them to retain functional properties close to the original WT structures.

Regions of a protein with high RMSF values often correlate with higher SASA values and vice versa [51]. Flexible regions tend to be more solvent-exposed because they can move more freely, increasing their surface area that interacts with the solvent [52]. In our study, the mutant complexes of D60Y and A14990Gfs exhibit lower RMSF values compared to the WT complexes, indicating increased stability and reduced flexibility. Additionally, these mutants show decreased SASA values, suggesting that their surface areas exposed to the solvent are also reduced. This combination of lower RMSF and SASA values implies that the variations lead to a more rigid and compact structure. The rigidity of proteins and their interactions possibly are affected by fluctuations in total intra- and intermolecular-H-bonds. Average number of H-bond present in D60Y mutant complex is 5 throughout the simulation as compared to WT (average HBN=3). Similarly A14990Gfs mutant complex show more HBN than WT strongly supporting the aforementioned rigidity and compactness of the complex.

Lu et al., (1998) employed steered molecular dynamics (SMD) to investigate the mechanical unfolding of Ig domains in TTN. The simulations have revealed that these domains unfold via a stepwise mechanism, where the sequential breaking of hydrogen bonds contributes to the protein’s elasticity [53]. Subsequently, Puchner et al., (2008) describe the effects of variations on TTN mechanics explaining specific variations can either destabilize the protein or enhance its mechanical resilience, providing insights into the molecular basis of certain myopathies [54]. Hence, it is evident that the variation within different domains of TTN either gives rise to a markedly rigid structure or generates unstable protein-protein complexes. Prolonged strong binding is observed in TCAP-binding D60Y and Myosin-binding A14990Gfs mutant that may cause loss of cellular dynamics, formation of protein aggregates, mis-localization and conformational stress within sarcomere. Proteins often need to move to different cellular compartments to function correctly. Strong binding could prevent such movement, leading to mis-localization and subsequent dysfunction. In contrast, transient unstable interactions, seen in myosin-binding R18842H, E24390Kfs, R22555H, E30120K and lamin-binding Q33877X possibly allow non-specific or promiscuous interactions leading to impaired signaling and regulatory mechanism.

## Conclusion

Our study establishes a framework for using combined genomic and *in-silico* methods to investigate the molecular basis of cardiomyopathies. We emphasize the utility of protein modeling and docking to understand the relations of protein structure and function. Since the large size of the TTN hinders conducting *in vivo* and *in vitro* experiments, therefore, computational methods are necessarily exploited to understand aberrant *TTN* functions. To validate the potential pathogenic contribution of the variations, more than one prediction algorithms are used in our study. Although bioinformatics serves as an ally of experimental procedures, we believe computational methods outlined in this article will be beneficial in creating a foundational framework to address variations in such large proteins. Further MD simulation offers deeper insights into the behaviour of molecules in a dynamic environment showing structural and functional properties of biological systems. Notably, high frequency of TTN variations in DCM patients and their clustering in the A-band region indicates their important role in myosin binding and contribution to the contractile function of the sarcomere, likely impairing the actomyosin mechano-signaling pathway by disrupting muscle contraction-relaxation, leading to heart-failure.

## Acknowledgment

We extend our heartfelt gratitude to the patients and their families for their invaluable participation in this study. Our sincere thanks go to Prof. D. Agrawal from the Department of Cardiothoracic & Vascular Surgery and Prof. Ashok Kumar from the Department of Pediatrics at IMS, BHU, Varanasi. Their encouragement and constant support were crucial in enrolling DCM patients for this research. We acknowledge Department of Biotechnology, Ministry of Science & Technology, Govt. of India for project grant (BT/PR12369/MED/12/678/2014) and Indian Council of Medical Research (ICMR) for Senior research fellowship to Ms. Amrita Mukhopadhyay.

## Author contributions

Bhagyalaxmi Mohapatra: Conceptualization, Supervision, Investigation,Validation, Resources, Writing-Review & editing, Amrita Mukhopadhyay: Data curation, Validation, Formal analysis, Visualization, Writing-Original draft preparation, Bharti Devi: Software, Formal analysis, Writing-review & editing, Anurag TK Baidya: Software, Writing-Review & Editing, Manohar lal Yadav: Software, Formal analysis, Rajnish Kumar: Resources, Writing-Reviewing and Editing.

## References

1. Giri P, Mukhopadhyay A, Gupta M, Mohapatra B. Dilated cardiomyopathy: a new insight into the rare but common cause of heart failure. Heart Failure Reviews. 2022 Mar;27(2):431–54. 10.1007/s10741-021-10125-6

2. Dec GW, Fuster V. Idiopathic dilated cardiomyopathy. N Engl J Med (1994);331:1564-75. 10.1056/NEJM199412083312307

3. Hershberger RE, Morales A, Siegfried JD. Clinical and genetic issues in dilated cardiomyopathy: a review for genetics professionals. Genetics in Medicine. (2010) Nov 1;12(11):655–67. 10.1097/GIM.0b013e3181f2481f

4. Dellefave L, McNally EM. The genetics of dilated cardiomyopathy. Curr Opin Cardiol (2010);25:198–204. 10.1097/HCO.0b013e328337ba52

5. Freiburg A, Trombitas K, Hell W, Cazorla O, Fougerousse F, Centner T, Kolmerer B, Witt C, Beckmann JS, Gregorio CC, Granzier H. Series of exon-skipping events in the elastic spring region of titin as the structural basis for myofibrillar elastic diversity. Circulation research. (2000) Jun 9;86(11):1114-21. 10.1161/01.res.86.11.1114

6. Hidalgo C, Granzier H. Tuning the molecular giant titin through phosphorylation: role in health and disease. Trends in cardiovascular medicine. (2013) Jul 1;23(5):165-71. 10.1016/j.tcm.2012.10.005

7. Bang ML, Centner T, Fornoff F, Geach AJ, Gotthardt M, McNabb M, Witt CC, Labeit D, Gregorio CC, Granzier H, Labeit S. The complete gene sequence of titin, expression of an unusual≈ 700-kDa titin isoform, and its interaction with obscurin identify a novel Z-line to I-band linking system. Circulation research. (2001) Nov 23;89(11):1065-72. 10.1161/hh2301.100981

8. Herman DS, Lam L, Taylor MR, Wang L, Teekakirikul P, Christodoulou D, Conner L, DePalma SR, McDonough B, Sparks E, Teodorescu DL. Truncations of titin causing dilated cardiomyopathy. New England Journal of Medicine. (2012) Feb 16;366(7):619-28. 10.1056/NEJMoa1110186

9. Li J, Shi L, Zhang K, Zhang Y, Hu S, Zhao T, Teng H, Li X, Jiang Y, Ji L, Sun Z. VarCards: an integrated genetic and clinical database for coding variants in the human genome. Nucleic acids research. (2018) Jan 4;46(D1):D1039-48. 10.1093/nar/gkx1039

10. Ashkenazy H, Abadi S, Martz E, Chay O, Mayrose I, Pupko T, et al. ConSurf 2016: an improved methodology to estimate and visualize evolutionary conservation in macromolecules. Nucleic Acids Res. (2016);44(W1):W344–50. 10.1093/nar/gkw408

11. McGuffin LJ, Bryson K, Jones DT. The PSIPRED protein structure prediction server. Bioinformatics. (2000);16(4):404–5. 10.1093/bioinformatics/16.4.404

12. Dominguez C, Boelens R, Bonvin AM. HADDOCK: a protein− protein docking approach based on biochemical or biophysical information. Journal of the American Chemical Society. 2003 Feb 19;125(7):1731–7. 10.1021/ja026939x

13. Bianco P, Nagy A, Kengyel A, Szatmári D, Mártonfalvi Z, Huber T, Kellermayer MS. Interaction forces between F-actin and titin PEVK domain measured with optical tweezers. Biophysical Journal. (2007) Sep 15;93(6):2102-9. 10.1529/biophysj.107.106153

14. Linke WA, Kulke M, Li H, Fujita-Becker S, Neagoe C, Manstein DJ, Gautel M, Fernandez JM. PEVK domain of titin: an entropic spring with actin-binding properties. Journal of structural biology. (2002) Jan 1;137(1-2):194-205. 10.1006/jsbi.2002.4468

15. Kulke M, Fujita-Becker S, Rostkova E, Neagoe C, Labeit D, Manstein DJ, Gautel M, Linke WA. Interaction between PEVK-titin and actin filaments: origin of a viscous force component in cardiac myofibrils. Circulation Research. (2001) Nov 9;89(10):874-81. 10.1161/hh2201.099453

16. Nagy A, Cacciafesta P, Grama L, Kengyel A, Málnási-Csizmadia A, Kellermayer MS. Differential actin binding along the PEVK domain of skeletal muscle titin. Journal of cell science. (2004) Nov 15;117(24):5781-9. 10.1242/jcs.01501

17. Gregorio CC, Trombitás K, Centner T, Kolmerer B, Stier G, Kunke K, Suzuki K, Obermayr F, Herrmann B, Granzier H, Sorimachi H. The NH2 terminus of titin spans the Z-disc: its interaction with a novel 19-kD ligand (T-cap) is required for sarcomeric integrity. The Journal of cell biology. (1998) Nov 16;143(4):1013-27. 10.1083/jcb.143.4.1013

18. Zou P, Pinotsis N, Lange S, Song YH, Popov A, Mavridis I, Mayans OM, Gautel M, Wilmanns M. Palindromic assembly of the giant muscle protein titin in the sarcomeric Z-disk. Nature. (2006) Jan 12;439(7073):229-33. 10.1038/nature04343

19. Pinotsis N, Petoukhov M, Lange S, Svergun D, Zou P, Gautel M, Wilmanns M. Evidence for a dimeric assembly of two titin/telethonin complexes induced by the telethonin C-terminus. Journal of structural biology. (2006) Aug 1;155(2):239-50. 10.1038/nature04343

20. Tharp CA, Haywood ME, Sbaizero O, Taylor MR, Mestroni L. The giant protein titin’s role in cardiomyopathy: genetic, transcriptional, and post-translational modifications of TTN and their contribution to cardiac disease. Frontiers in Physiology. (2019) Nov 28;10:1436. 10.3389/fphys.2019.01436

21. Linke WA. Sense and stretchability: the role of titin and titin-associated proteins in myocardial stress-sensing and mechanical dysfunction. Cardiovascular research. (2008) Mar 1;77(4):637-48. 10.1016/j.cardiores.2007.03.029

22. Blair E, Redwood C, de Jesus Oliveira M, Moolman-Smook JC, Brink P, Corfield VA, Östman-Smith I, Watkins H. Mutations of the light meromyosin domain of the β-myosin heavy chain rod in hypertrophic cardiomyopathy. Circulation research. (2002) Feb 22;90(3):263-9. 10.1161/hh0302.104532

23. Sohn RL, Vikstrom KL, Strauss M, Cohen C, Szent-Gyorgyi AG, Leinwand LA. A 29 residue region of the sarcomeric myosin rod is necessary for filament formation. Journal of molecular biology. (1997) Feb 21;266(2):317-30. 10.1006/jmbi.1996.0790

24. Abraham MJ, Murtola T, Schulz R, Páll S, Smith JC, Hess B, et al. GROMACS: High performance molecular simulations through multi-level parallelism from laptops to supercomputers. SoftwareX. (2015);1:19–25. 10.1016/j.softx.2015.06.001

25. Jo S, Cheng X, Lee J, Kim S, Park SJ, Patel DS, et al. CHARMM-GUI 10 years for biomolecular modeling and simulation. Journal of computational chemistry. (2017);38(15):1114–24. 10.1002/jcc.24660

26. Lee J, Cheng X, Jo S, MacKerell AD, Klauda JB, Im W. CHARMM-GUI input generator for NAMD, GROMACS, AMBER, OpenMM, and CHARMM/OpenMM simulations using the CHARMM36 additive force field. Biophysical journal. (2016);110(3):641a. 10.1021/acs.jctc.5b00935

27. Huang J, Rauscher S, Nawrocki G, Ran T, Feig M, De Groot BL, et al. CHARMM36m: an improved force field for folded and intrinsically disordered proteins. Nature methods. (2017);14(1):71–3. 10.1038/nmeth.4067

28. Morozova TI, García NA, Barrat J-L. Temperature dependence of thermodynamic, dynamical, and dielectric properties of water models. The Journal of Chemical Physics. (2022);156(12). 10.1063/5.0079003

29. Martoňák R, Laio A, Parrinello M. Predicting crystal structures: the Parrinello-Rahman method revisited. Physical review letters. (2003);90(7):075503. 10.1103/PhysRevLett.90.075503

30. Rodrigues CHM, Pires DE V, Ascher DB. DynaMut2: Assessing changes in stability and flexibility upon single and multiple point missense mutations. ProteinSci. (2021);30(1):60–9. 10.1002/pro.3942

31. Wainreb G, Wolf L, Ashkenazy H, Dehouck Y, Ben-Tal N. Protein stability: a single recorded mutation aids in predicting the effects of other mutations in the same amino acid site. Bioinformatics. (2011) Dec 1;27(23):3286-92. 10.1093/bioinformatics/btr576

32. Parthiban V, Gromiha MM, Schomburg D. CUPSAT: prediction of protein stability upon point mutations. Nucleic Acids Res. (2006);34(suppl_2):W239–42. 10.1093/nar/gkl190

33. Pires DE V, Ascher DB, Blundell TL. DUET: a server for predicting effects of mutations on protein stability using an integrated computational approach. Nucleic Acids Res. (2014);42(W1):W314–9. 10.1093/nar/gku411

34. Capriotti E, Fariselli P, Casadio R. I-Mutant2. 0: predicting stability changes upon mutation from the protein sequence or structure. Nucleic acids research. (2005) Jul 1;33(suppl_2):W306–10. 10.1093/nar/gki375

35. Roy S, Maheshwari N, Chauhan R, Sen NK, Sharma A. Structure prediction and functional characterization of secondary metabolite proteins of Ocimum. Bioinformation. (2011);6(8):315. 10.6026/97320630006315

36. Laskowski RA, Thornton JM. PDBsum extras: SARS-Cov-2 and AlphaFold models. Protein science. (2022) Jan;31(1):283-9. 10.1002/pro.4238

37. Begay RL, Graw S, Sinagra G, Merlo M, Slavov D, Gowan K, Jones KL, Barbati G, Spezzacatene A, Brun F, Di Lenarda A. Role of titin missense variants in dilated cardiomyopathy. Journal of the American Heart Association. (2015) Nov 13;4(11):e002645. 10.1161/JAHA.115.002645

38. Haas J, Frese KS, Peil B, Kloos W, Keller A, Nietsch R, Feng Z, Müller S, Kayvanpour E, Vogel B, Sedaghat-Hamedani F. Atlas of the clinical genetics of human dilated cardiomyopathy. European heart journal. 2015 May 7;36(18):1123–35. 10.1093/eurheartj/ehu301

39. Kellermayer D, Smith JE, Granzier H. Titin mutations and muscle disease. Pflügers Archiv-European Journal of Physiology. (2019) May 1;471:673–82. 10.1007/s00424-019-02272-5

40. Franaszczyk M, Chmielewski P, Truszkowska G, Stawinski P, Michalak E, Rydzanicz M, Sobieszczanska-Malek M, Pollak A, Szczygieł J, Kosinska J, Parulski A. Titin truncating variants in dilated cardiomyopathy–prevalence and genotype-phenotype correlations. PLoS One. (2017) Jan 3;12(1):e0169007. 10.1371/journal.pone.0169007

41. Pugh TJ, Kelly MA, Gowrisankar S, Hynes E, Seidman MA, Baxter SM, Bowser M, Harrison B, Aaron D, Mahanta LM, Lakdawala NK. The landscape of genetic variation in dilated cardiomyopathy as surveyed by clinical DNA sequencing. Genetics in Medicine. (2014) Aug;16(8):601-8. 10.1038/gim.2013.204

42. Muhle-Goll C, Habeck M, Cazorla O, Nilges M, Labeit S, Granzier H. Structural and functional studies of titin’s fn3 modules reveal conserved surface patterns and binding to myosin S1-a possible role in the frank-starling mechanism of the heart. Journal of molecular biology. (2001) Oct 19;313(2):431-47. 10.1006/jmbi.2001.5017

43. Li S, Dohlman HG. Evolutionary conservation of sequence motifs at sites of protein modification. Journal of Biological Chemistry. 2023 May 1;299(5). 10.1016/j.jbc.2023.104617

44. Nooren IM, Thornton JM. Diversity of protein–protein interactions. The EMBO journal. (2003) Jul 15;22(14):3486-92. 10.1093/emboj/cdg359

45. Petukh M, Kucukkal TG, Alexov E. On human disease-causing amino acid variants: Statistical study of sequence and structural patterns. Human mutation. (2015) May;36(5):524-34. 10.1002/humu.22770

46. Tokmakov AA, Kurotani A, Sato KI. Protein pI and intracellular localization. Frontiers in Molecular Biosciences. (2021) Nov 29;8:775736. 10.3389/fmolb.2021.775736

47. Lee EH, Hsin J, Sotomayor M, Comellas G, Schulten K. Discovery through the computational microscope. Structure. (2009) Oct 14;17(10):1295-306. 10.1016/j.str.2009.09.001

48. Suganthi M, Sowmya H, Manjunathan J, Ramasamy P, Thiruvengadam M, Varadharajan V, Venkidasamy B, Senthilkumar P. Homology modeling and protein-protein interaction studies of GAPDH from Helopeltis theivora and chitinase from Pseudomonas fluorescens to control infection in tea (Camellia sinensis (L.) O. Kuntze) plants. Plant Stress. (2024) Mar 1;11:100377. 10.1016/j.stress.2024.100377

49. Wang Y, Parmar S, Schneekloth JS, Tiwary P. Interrogating RNA–small molecule interactions with structure probing and artificial intelligence-augmented molecular simulations. ACS Central Science. (2022) May 16;8(6):741-8. 10.1021/acscentsci.2c00149

50. Vassetti D, Civalleri B, Labat F. Analytical calculation of the solvent-accessible surface area and its nuclear gradients by stereographic projection: A general approach for molecules, polymers, nanotubes, helices, and surfaces. Journal of Computational Chemistry. (2020) Jun 5;41(15):1464-79. 10.1002/jcc.26191

51. Boroujeni MB, Dastjerdeh MS, Shokrgozar M, Rahimi H, Omidinia E. Computational driven molecular dynamics simulation of keratinocyte growth factor behavior at different pH conditions. Informatics in Medicine Unlocked. 2021 Jan 1;23:100514. 10.1016/j.imu.2021.100514

52. Ranjan P, Das P. An inclusive study of deleterious missense PAX9 variants using user-friendly tools reveals structural, functional alterations, as well as potential therapeutic targets. International Journal of Biological Macromolecules. 2023 Apr 1;233:123375. 10.1016/j.ijbiomac.2023.123375

53. Lu H, Isralewitz B, Krammer A, Vogel V, Schulten K. Unfolding of titin immunoglobulin domains by steered molecular dynamics simulation. Biophysical journal. (1998) Aug 1;75(2):662-71. 10.1016/S0006-3495(98)77556-3

54. Puchner EM, Alexandrovich A, Kho AL, Hensen U, Schäfer LV, Brandmeier B, Gräter F, Grubmüller H, Gaub HE, Gautel M. Mechanoenzymatics of titin kinase. Proceedings of the National Academy of Sciences. (2008) Sep 9;105(36):13385-90. 10.1073/pnas.0805034105

