## Supplementary material for "Implication of *TITIN* Variations in Dilated Cardiomyopathy: Integrating Whole Exome Sequencing With Molecular Dynamics Simulation Study"

***Supplementary data***

***Supplementary Table 1: List of identified genetic variants in TTN***

| Patient Id  (LVEF) | cDNA Change | Amino-Acid Change  (dbSNP) | Type Of Mutation | *TTN*  Domains |
| --- | --- | --- | --- | --- |
| FDCM1  (LVEF=16%) | c.27362C>T | R9121Q  (rs72648998) | MISSENSE | Ig like 74  (I-band) |
|  | c.19910G>A | A6637V  (rs17355446) | MISSENSE | Ig like 48  (I-band) |
|  | c.24323C>T | S8108N  (rs11888217) | MISSENSE | Ig like 61  (I-band) |
|  | c.9781C>T | V3261M  (rs72648937) | MISSENSE | Ig like 19  (I-band) |
|  | c.9461T>C | K3154R  (rs4893853) | MISSENSE | Ig like 18-19  (I-band) |
| FDCM2  (LVEF 50%) | c.30913G>A | G10305R  (rs2244492) | MISSENSE | PEVK 4  (I-band) |
|  | c.30613A>G | I10205V  (rs2042995) | MISSENSE | Basic and acidic residues (I-band) |
|  | c.28202T>C | I9401T  (rs4893852) | MISSENSE | Ig like 77  (I-band) |
|  | c.25730C>T | P8577L  (rs13398235) | MISSENSE | Ig like 68  (I-band) |
|  | c.25457A>G | N8486S  (rs12693164) | MISSENSE | Ig like 67  (I-band) |
|  | c.23480A>C | E7827A  (rs16866465) | MISSENSE | Ig like 60  (I-band) |
|  | c.21835G>C | D7279H  (rs72648970) | MISSENSE | Ig like 54  (I-band) |
|  | c.21433G>C | D7145H  (rs12693166) | MISSENSE | Ig like 53  (I-band) |
|  | c.18350G>A | S6117N  (rs11888217) | MISSENSE | Ig like 42  (I-band) |
|  | c.13747G>A | A4583T  (rs72648923) | MISSENSE | Ig like 26  (I-band) |
|  | c.10471C>T | P3491S  (rs2627037) | MISSENSE | Coiled coil  (I-band) |
|  | c.10256G>A | S3419N  (rs2291310) | MISSENSE | Ig like 20  (I-band) |
|  | c.7174G>A | G2392S  (rs4894048) | MISSENSE | Ig like 13  (I-band) |
|  | c.4322G>C | R1441P  (rs72647876) | MISSENSE | ZIS 5  (Z-disc bound) |
|  | c.3884C>T | S1295L  (rs1552280) | MISSENSE | Ig like 5  (Z-disc bound) |
|  | c.3601A>G | K1201E  (rs10497520) | MISSENSE | Ig like 4-5  (Z-disc bound) |
|  | c.982C>T | R328C  (rs16866538) | MISSENSE | Before disordered  (Z-disc bound) |
|  | c.178G>T | D60Y  (rs35683768) | MISSENSE | Ig like 1  (Z-disc bound) |
| FDCM3  (LVEF=13%) | c.89122C>T | R29708C  (rs727503549) | MISSENSE | FnIII 116  (A-band) |
|  | c.67664G>A | R22555H  (rs200317412) | MISSENSE | FnIII63  (A-band) |
|  | c.21332-11del | NA  (rs778596216) | INTRONIC | - |
|  | c.20828C>A | S6943Y  (rs187925021) | MISSENSE | Ig like 51  (I-band) |
|  | c.20206A>C | T6736P  (rs727504741) | MISSENSE | Ig like 49  (I-band) |
| FDCM4  (LVEF=35%) | c.101696T>C | I33899T  (rs55880440) | MISSENSE | Ig like 149-150  (M-band) |
|  | c.83587G>A | D27863N  (rs376679796) | MISSENSE | Ig like 130  (A-band) |
|  | c.44969AG>A | A14990GfsTer4  (Novel) | FRAME-SHIFT DEL | FnIII8  (A-band) |
|  | c.48584G>A | R16195H  (rs373526624) | MISSENSE | FnIII 17  (A-band) |
|  | c.47068G>C | E15690Q  (rs547986881) | MISSENSE | Ig like 98  (A-band) |
| FDCM5  (LVEF=20%) | 93371C>G | A31124G  (rs72648273) | MISSENSE | Ig like 139  (A-band) |
|  | c.41560G>A | A13854T  (rs537428006) | MISSENSE | Ig like 94-95  (I-band) |
|  | c.28861A>T | T9621S  (rs72650006) | MISSENSE | Before Ig like 78 (I-band) |
|  | c.168-8C>T | NA  (rs59415652) | INTRONIC | - |
| SDCM1  (LVEF=62%) | c.36655T>G | L12219V  (rs12994774) | MISSENSE | Basic and acid residues  (I-band) |
| SDCM2  (LVEF=26%) | c.100257G>C | E33419D  (rs56308529) | MISSENSE | After ig like 146  (M-band) |
|  | c.22072G>T | D7358Y  (rs552951988) | MISSENSE | Ig like 55  (I-band) |
| SDCM3  (LVEF=21%) | c.4480+6C>T | NA  (rs719201) | INTRONIC | - |
|  | c.59G>A | NA  (rs72629795) | 3’UTR | - |
|  | c.101752G>C | E33918Q  (rs199632397) | MISSENSE | Ig like 149-150  (M-band) |
|  | c.78797G>A | S26266N  (rs759630792) | MISSENSE | FnIII 91  (A-band) |
|  | c.61871C>T | A20624V  (rs12463674) | MISSENSE | FnIII49-50  (A-band) |
|  | c.61463G>A | R20488H  (rs12464787) | MISSENSE | FnIII 48  (A-band) |
|  | c.60852C>T | S20284S  (rs4145333) | SILENT | Ig like 111  (A-band) |
|  | c.11284A>G | I3762V  (rs750836266) | MISSENSE | After ig like 22  (I-band) |
| SDCM4  (LVEF=15%) | c.81760G>A | V27254M  (rs201290358) | MISSENSE | FnIII 98  (A-band) |
|  | c.60224C>T | S20075L  (rs13021201) | MISSENSE | FnIII 45  (A-band) |
|  | c.21139C>T | R7047W  (rs397517500) | MISSENSE | Ig like 52  (I-band) |
| SDCM5  (LVEF=18-20%) | c.102166G>C | E34056Q  (rs199531140) | MISSENSE | Ig-like 150-151  (M-band) |
|  | c.71420G>A | S23807N  (rs3813243) | MISSENSE | FnIII 73  (A-band) |
|  | c.65174T>C | V21725A  (rs372782502) | MISSENSE | Ig-like 114  (A-band) |
|  | c.60169C>T | R20057C  (rs72646861) | MISSENSE | Fn III 45  (A-band) |
|  | c.56525G>A | R18842H  (Novel) | MISSENSE | Ig-like 107  (A-band) |
|  | c.46559C>T | A15520V  (rs16866412) | MISSENSE | FnIII 13  (A-band) |
|  | c.43858G>A | D14620N  (rs764203042) | MISSENSE | Ig-like 96  (A-band) |
|  | c.35183C>G | P11728R  (rs72650066) | MISSENSE | PEVK 23  (I-band) |
|  | c.20093C>T | A6698V  (rs72648960) | MISSENSE | Ig-like 48  (I-band) |
|  | c.6359G>T | R2120L  (rs141142920) | MISSENSE | Ig-like 10  (I-band) |
|  | c.78090_78094del CTTTCT>C | E24390KfsTer15  (Novel) | FRAMESHIFT-DEL | Fn III 77  (A-band) |
|  | c.105582C>T | S33553S  (rs3829749) | SILENT | Ig like 147  (M-band) |
|  | c.52821T>C | D15966D  (rs2303831) | SILENT | FnIII 16  (A-band) |
|  | c.49758T>C | Y14945Y  (rs72677247) | SILENT | FnIII 8  (A-band) |
|  | c.49731T>C | H14936H  (rs2115558) | SILENT | FnIII 8  (A-band) |
|  | c.48996G>A | E14691E  (rs72677244) | SILENT | Ig like 96  (A-band) |
|  | c.43596T>C | N12891N  (rs16866423) | SILENT | Ig like 85-86  (M-band) |
|  | c.36625G>T  NM_001267550 | V12209L  (rs72650053) | MISSENSE | Ig-like domain  (I-band) |
| SDCM6  (LVEF=35%) | c.79525T>C | Y26509H  (rs397517727) | MISSENSE | FnIII92  (A-band) |
|  | c.60109G>A | V20037I  (rs570562262) | MISSENSE | FnIII45  (A-band) |
|  | c.49182G>A | A16394A  (rs371155050) | SILENT | Ig like 100  (A-band) |
|  | c.46185A>G | A15395A  (rs748295772) | SILENT | Ig like 97  (A-band) |
|  | c.27060C>T | D9020D  (rs757848062) | SILENT | Ig like 73  (I-band) |
| SDCM7  (LVEF=28%) | c.102344T>C | V34115A  (rs16866378) | MISSENSE | Ig like 151  (M-band) |
|  | c.100859C>T | P33620L  (rs16866380) | MISSENSE | Before Ig like 148 (M-band) |
|  | c.97910G>T | G32637V  (rs3731752) | MISSENSE | Ig like 144  (M-band) |
|  | c.90358G>A | E30120K  (rs727505342) | MISSENSE | fnIII 119  (A-band) |
|  | c.74139T>A | G24713G  (rs3731744) | SILENT | FnIII 79  (A-band) |
|  | c.56301G>A | V18767V  (rs566188777) | SILENT | FnIII 36  (A-band) |
| SDCM8  (LVEF=28%) | c.101629C>T | Q33877X  (Novel) | NONSENSE | Ig like 149-150  (M-band) |
|  | c.38055C>T | Y12685Y  (rs369959066) | SILENT | Ig like 84-85  (I-band) |
|  | c.22902C>A | A7634A  (rs201766927) | SILENT | Ig like 58  (I-band) |
|  | c.3695A>T | D1232V  (rs576019235) | MISSENSE | Ig like 4-5  (Z-disc bound) |
| SDCM9  (LVEF=21%) | c.91257T>C | I30419I  (rs572401798) | SILENT | Ig like 137  (A-band) |
|  | c.86147C>T | T28716I  (rs754537852) | MISSENSE | FnIII 109  (A-band) |
|  | c.67182T>C | F22394F  (rs397517691) | SILENT | Ig 116  (A-band) |
|  | c.57107T>C | I19036T  (rs558670891) | MISSENSE | FnIII 38  (A-band) |
|  | c.5875T>A | F1959I  (rs558670891) | MISSENSE | After ig like 9  (Z-disc bound) |
| SDCM10  (LVEF=34%) | c.77462C>T | T25821M  (rs55933739) | MISSENSE | FnIII 87  (A-band) |
|  | c.52797A>G: | R17599R  (rs574980683) | SILENT | FnIII 27  (A-band) |
|  | c.52796G>A | R17599Q  (rs535940583) | MISSENSE | FnIII 27  (A-band) |

*Abbreviations: FDCM, Familial Dilated Cardiomyopathy; SDCM, Sporadic Dilated Cardiomyopathy; rs-ID,*

*Reference SNP cluster ID; LVEF, Left-ventricular ejection fraction; TTN, Titin; Ig-like, Immunoglobulin-like,*

*FnIII-Fibronectin type III.*

***Supplementary Table 2: List of the genetic variations in TTN and their frequency in different databases***

| Amino acid (AA)  position | dbSNP | ExAc | gnomAD | 1000Genomes | Genome Asia | IndiGenomes | Index-db |
| --- | --- | --- | --- | --- | --- | --- | --- |
| D60Y | rs35683768 | 0.013900 | 0.013200 | 0.028953 | NR | 0.0103 | 0.00571 |
| R29708C | rs727503549 | 0 | 0.000089 | NR | NR | 0 | NR |
| R22555H | rs200317412 | 0 | 0.000247 | 0 | NR | 0 | NR |
| A14990GfsTer4 | Novel  (SCV3836535) | NR | NR | NR | NR | NR | NR |
| R16195H | rs373526624 | 0 | 0.000723 | 0 | NR | 0.0039 | 0.00286 |
| E15690Q | rs547986881 | 0 | 0.000698 | 0 | NR | 0.0044 | 0.00571 |
| A31124G | rs72648273 | 0 | 0.003032 | 0 | NR | 0.0005 | NR |
| R20488H | rs187257105 | 0.000165 | 0.000175 | NR | NR | 0.0005 | NR |
| V27254M | rs201290358 | 0.001749 | 0.001525 | 0.002995 | NR | 0.0103 | 0.0171 |
| S20075L | rs13021201 | 0.025638 | 0.024714 | 0.010782 | NR | 0.0186 | 0.00286 |
| S23807N | rs3813243 | 0.041270 | 0.041282 | 0.079273 | NR | 0.0463 | 0.05143 |
| E24390KfsTer15 | Novel  (SCV00401519) | NR | NR | NR | NR | NR | NR |
| A15520V | rs16866412 | 0.034652 | 0.034005 | 0.072684 | NR | 0.0591 | 0.05143 |
| R18842H | Novel  (SCV00401518) | 0.000018 | 0.000009 | NR | NR | NR | NR |
| R20057C | rs72646861 | 0.008826 | 0.008245 | 0.012979 | NR | 0.0328 | 0.04286 |
| P11728R | rs72650066 | 0.030576 | 0.018376 | 0.039337 | NR | NR | 0.04000 |
| Y26509H | rs397517727 | 0.000102 | 0.000091 | 0.000399 | NR | NR | 0.00286 |
| E30120K | rs727505342 | 0.000008 | 0.000012 | NR | NR | NR | NR |
| Q33877X | Novel  (SCV00256798) | NR | NR | NR | NR | NR | NR |
| I19036T | rs558670891 | 0.000041 | 0.000049 | 0.000200 | NR | 0.0005 | NR |
| T25821M | rs55933739 | 0.000066 | 0.000061 | NR | NR | NR | NR |

*Abbreviations: FDCM, Familial Dilated Cardiomyopathy; SDCM, Sporadic Dilated Cardiomyopathy; NR, Not Reported, rs-ID, Reference SNP cluster ID; ExAc, Exome Aggregation Consortium; gnomAD, Genome Aggregation Database; SCV, Clinical Variantion database accession numbers of the submitted novel variants*

***Supplementary Table 3: Prediction of docking Scores using HADDOCK tool***

| Complex Name | HADDOCK Score | Van Der Waals Energy (kcal/mol) | Electrostatic Energy  (kcal/mol) | Buried Surface Area  (kcal/mol) | Z-Score |
| --- | --- | --- | --- | --- | --- |
| Ig-Like1_Wild-type | 70.2 +/- 5.1 | -47.7 +/- 4.2 | -106.4 +/- 20.7 | 1481.7 +/- 117.4 | -2.1 |
| Ig-Like1_Mutant | 40.5 +/- 11.1 | -57.9 +/- 4.0 | -68.2 +/- 13.6 | 1622.0 +/- 64.7 | -1.8 |
| Ig-Like107_ Wild-type | -44.3 +/- 3.4 | -23.3 +/- 8.9 | -232.5 +/- 34.0 | 1097.5 +/- 36.7 | -1.6 |
| Ig-Like107_ Mutant | -60.8 +/- 1.7 | -24.9 +/- 4.7 | -289.1 +/- 29.6 | 1192.0 +/- 82.5 | -1.8 |
| Ig-Like149_ Wild-type | -11.3 +/- 5.5 | -49.2 +/- 2.3 | -201.6 +/- 10.7 | 1477.2 +/- 71.3 | -1.6 |
| Ig-Like149_ Mutant | -9.6 +/- 9.0 | -47.8 +/- 2.5 | -206.3 +/- 28.8 | 1388.1 +/- 172.7 | -2.1 |
| Fniii8_ Wild-type | -65.5 +/- 3.5 | -19.7 +/- 2.6 | -310.9 +/- 6.4 | 929.6 +/- 29.1 | -2.6 |
| Fniii8_ Mutant | -37.8 +/- 1.9 | -19.2 +/- 1.2 | -133.8 +/- 10.2 | 641.2 +/- 9.2 | -1.5 |
| Fniii63_ Wild-type | -50.7 +/- 2.4 | -26.5 +/- 2.5 | -148.4 +/- 17.4 | 771.7 +/- 54.9 | -0.6 |
| Fniii63_ Mutant | -43.2 +/- 2.1 | -28.2 +/- 4.0 | -71.9 +/- 16.0 | 940.2 +/- 29.6 | -0.6 |
| Fniii77_ Wild-type | -41.4 +/- 4.8 | -18.4 +/- 7.2 | -174.4 +/- 15.0 | 928.7 +/- 113.7 | -2 |
| Fniii77_ Mutant | -49.5 +/- 3.0 | -16.6 +/- 4.8 | -209.7 +/- 19.0 | 1032.5 +/- 60.5 | -1.8 |
| Fniii119_ Wild-type | -45.2 +/- 0.9 | -15.9 +/- 3.6 | -165.1 +/- 21.4 | 754.8 +/- 44.1 | -1.9 |
| Fniii119_ Mutant | -54.5 +/- 2.5 | -16.9 +/- 3.3 | -169.0 +/- 20.8 | 894.8 +/- 62.6 | -1.2 |

***Supplementary Figure 1: Phylogenetic Conservation of seven damaging variants across different species***

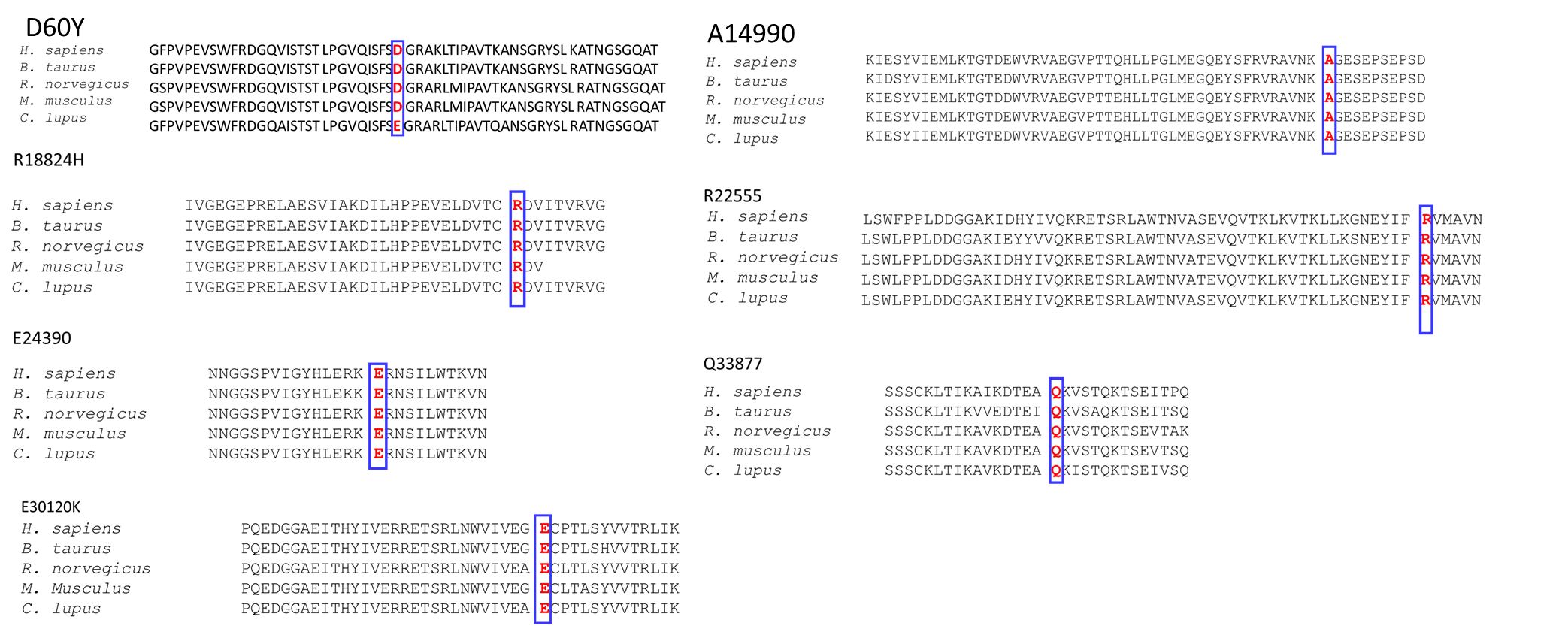

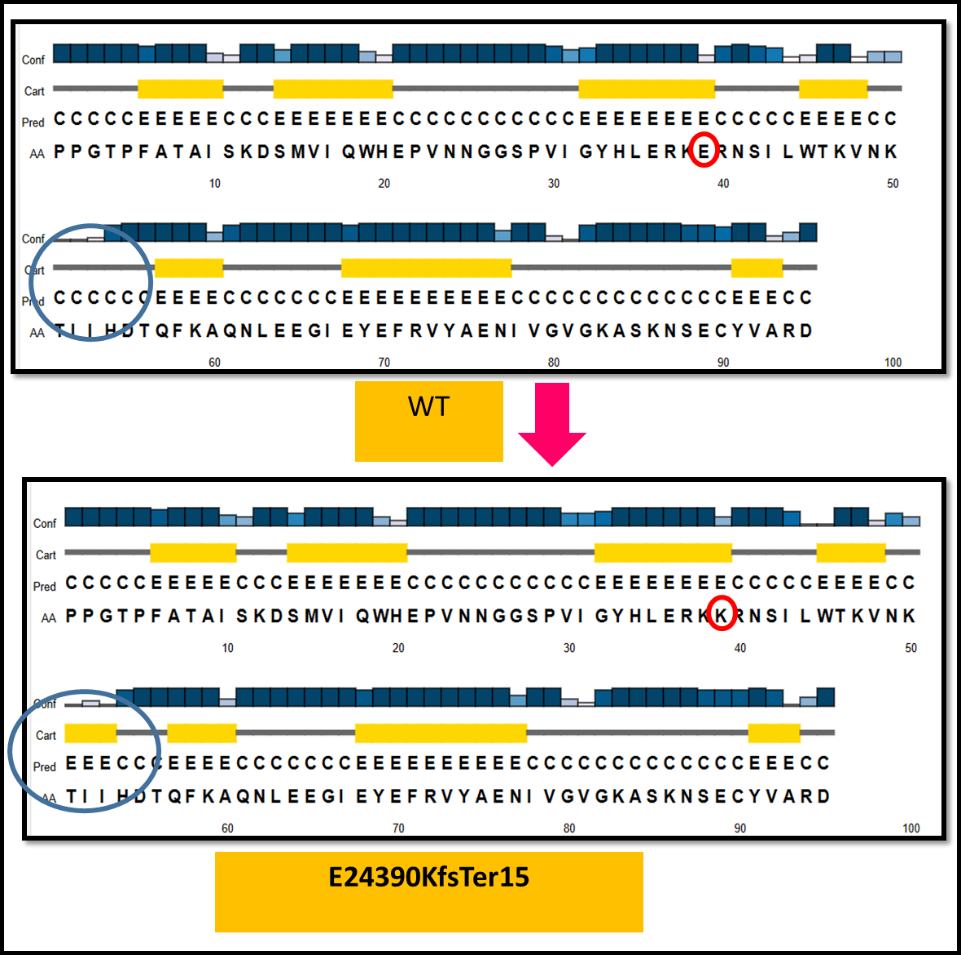

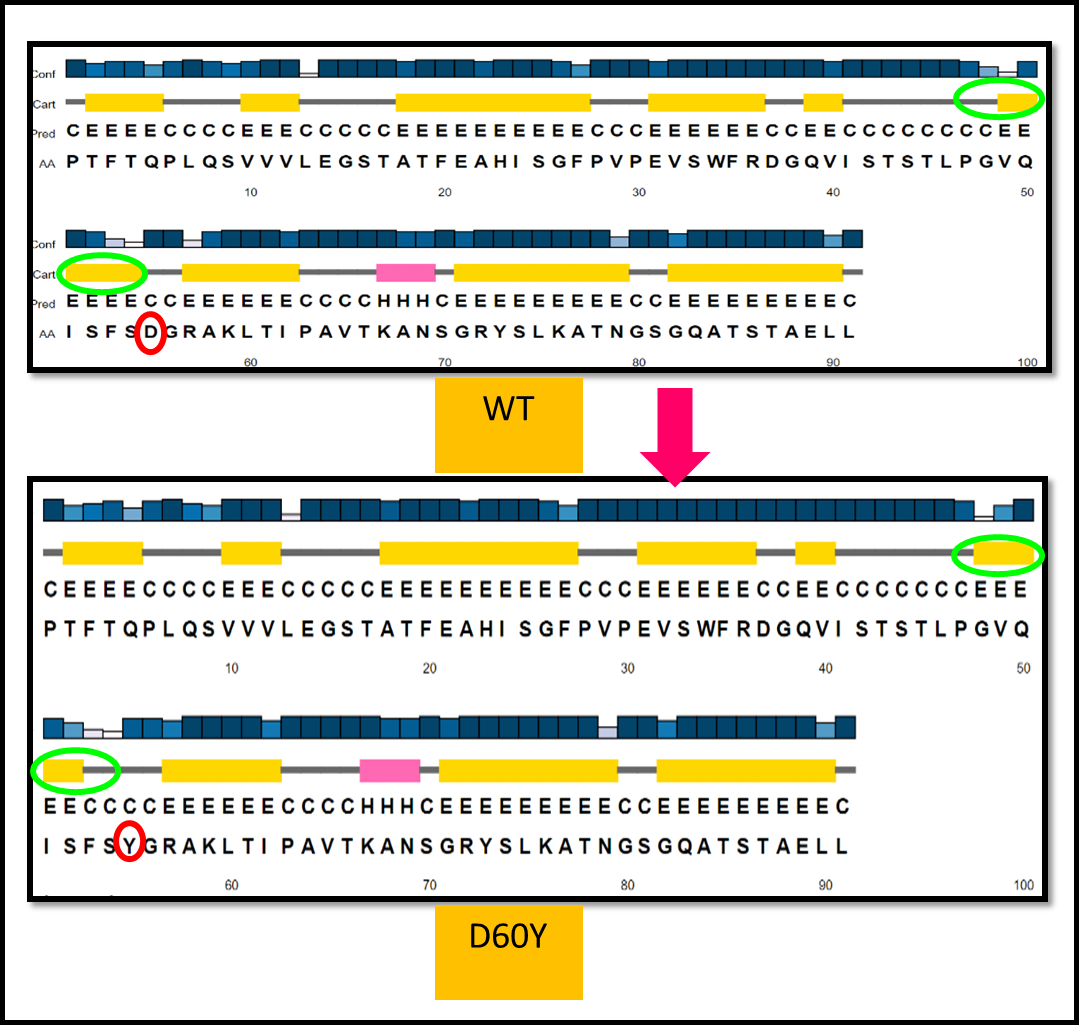

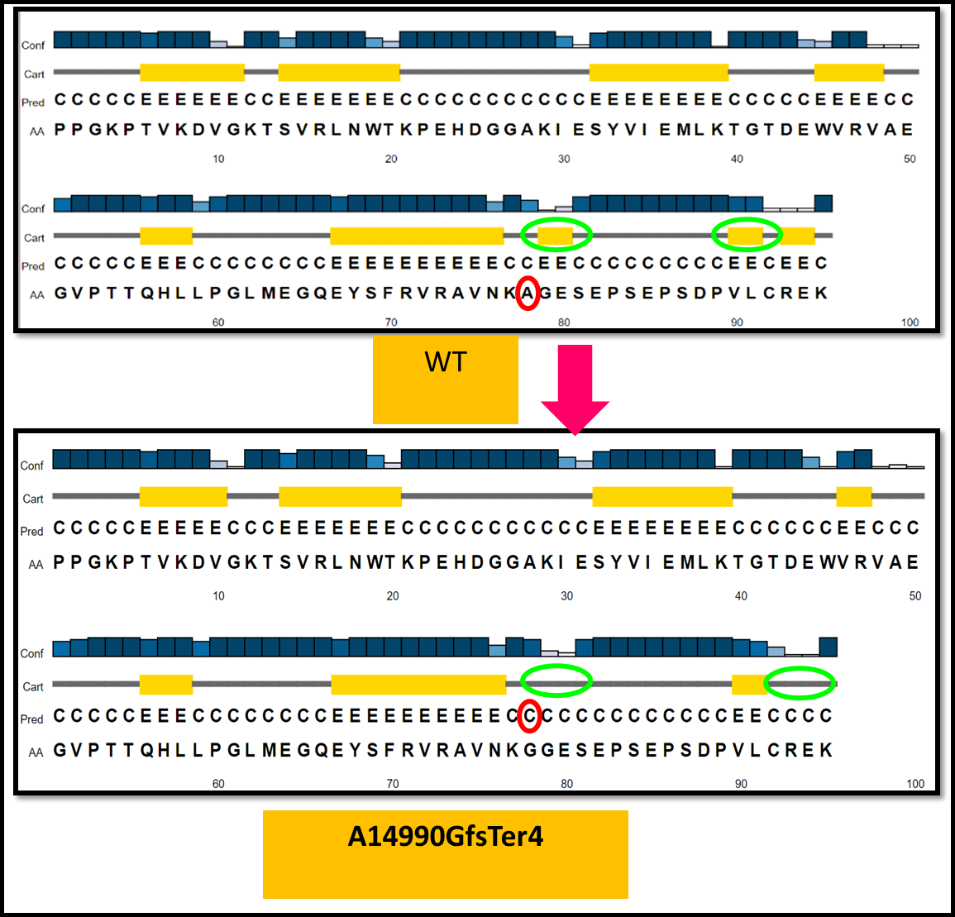

B

A

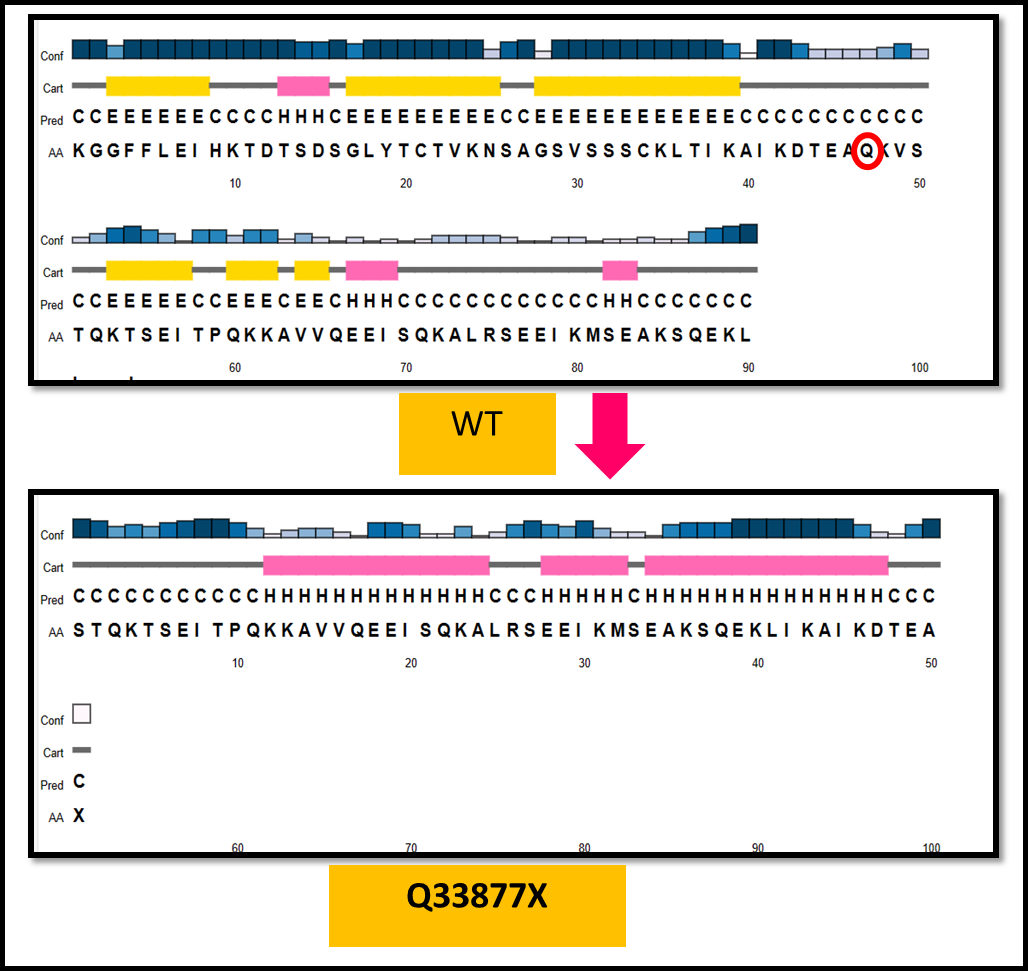

D

C

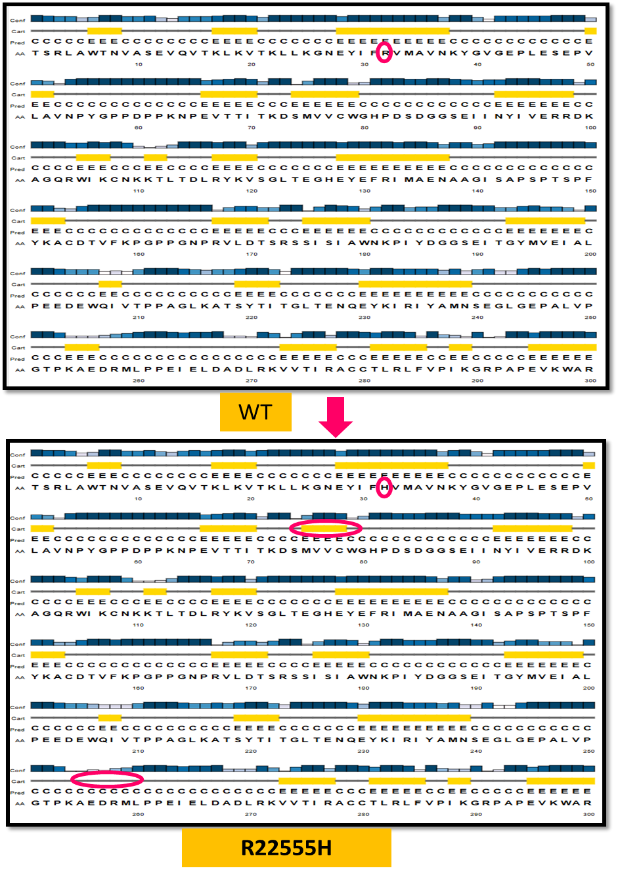

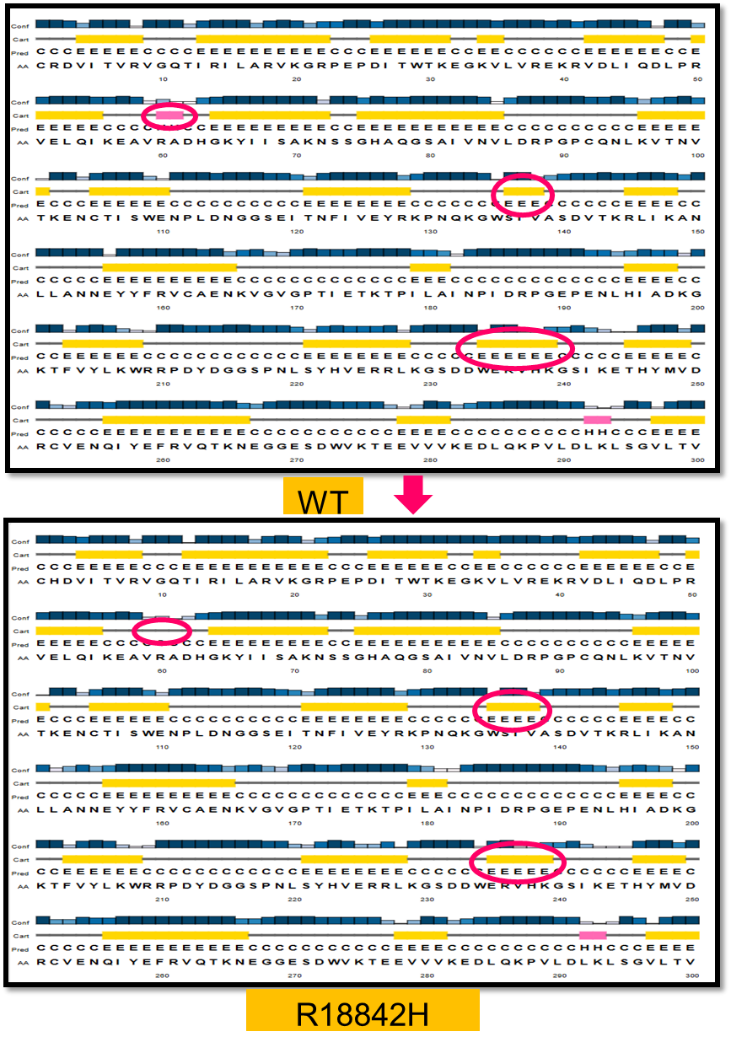

E

F

***Supplementary Figure 2:*** *Secondary structure prediction of TTN variants by PsiPred tool. Major changes in D60Y, A14990Gfs, E24390Kfs and Q33877X variants are shown here. In D60Y, the beta-strand was getting shorter upstream of the mutation that may disrupt protein’s functional capability; In A14990Gfs and E24390Kfs variant, beta strands that are present downstream of the mutation site was completely abolished that suspect functionally abnormal protein-protein interactions. Moreover in the nonsense mutation (Q33877X), the whole protein frame was distorted. In R18842H, one alpha-helix and two beta-strands are deformed downstream of the mutation. Similarly, in R22555H and in E30120K, majorly beta-strands are getting affected.*

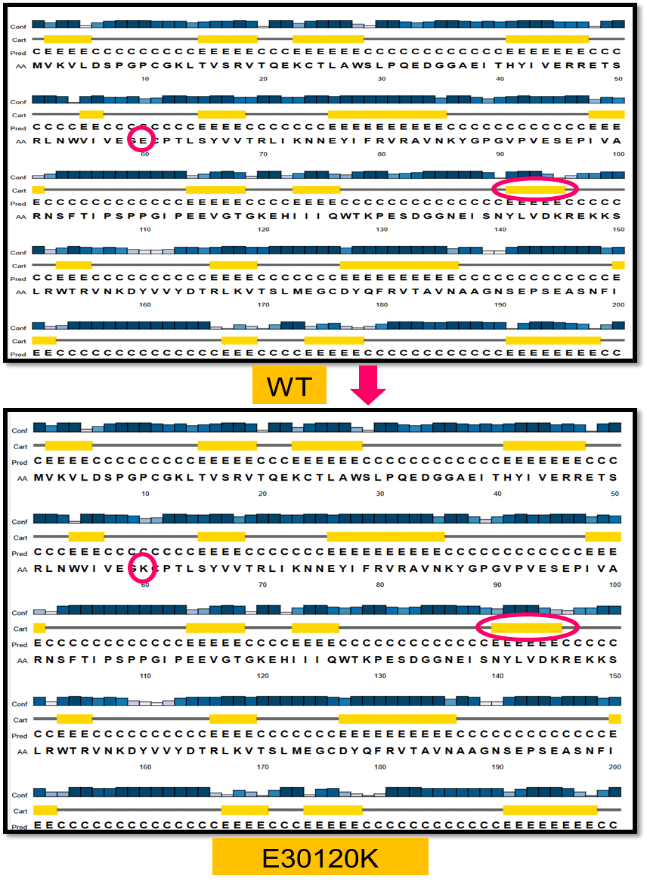

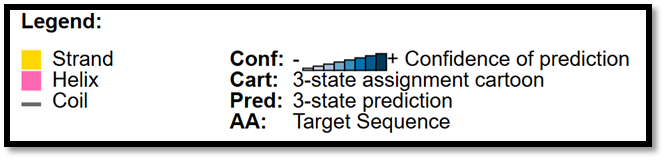

G

***Supplementary Figure 3A:*** *Representation of Sequence coverage plots that shows the number of homologs identified across the representative sequences are very high and denotes a highly conservative motif of TTN structure*

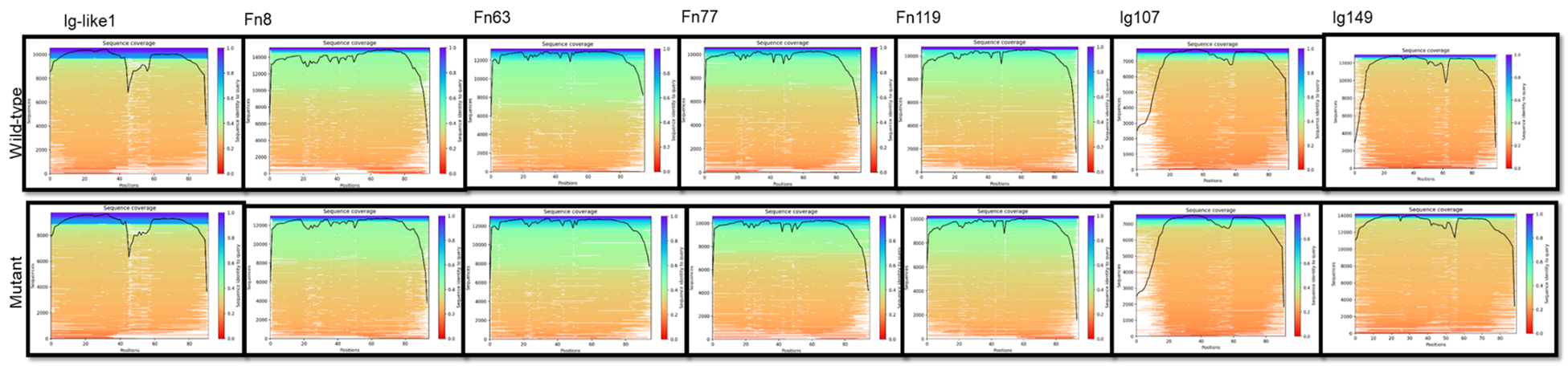

***Supplementary Figure 3B:*** *PAE plots of TTN wild-type and Mutant domains;* *It is displayed as an image for each of the structure predictions. It*

*indicates an expected positional error at residue x if the predicted and actual structures are aligned on residue y (using Cα, N, and C atoms).*

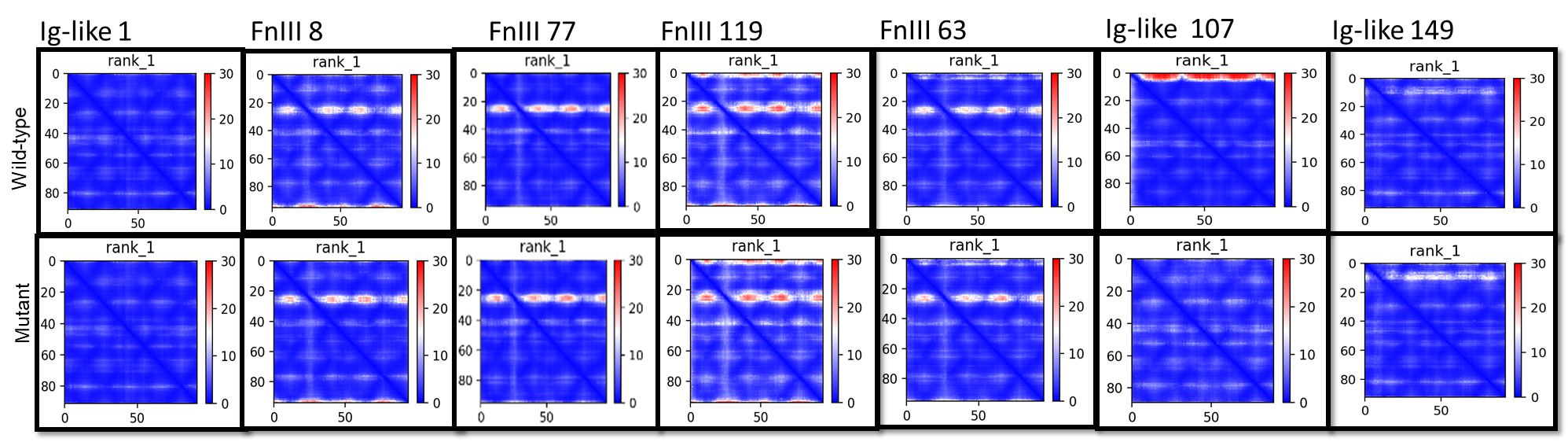

***Supplementary Figure 4: Analysis of Ramachandran Plots for Wild-Type and Mutant Domains of TTN***

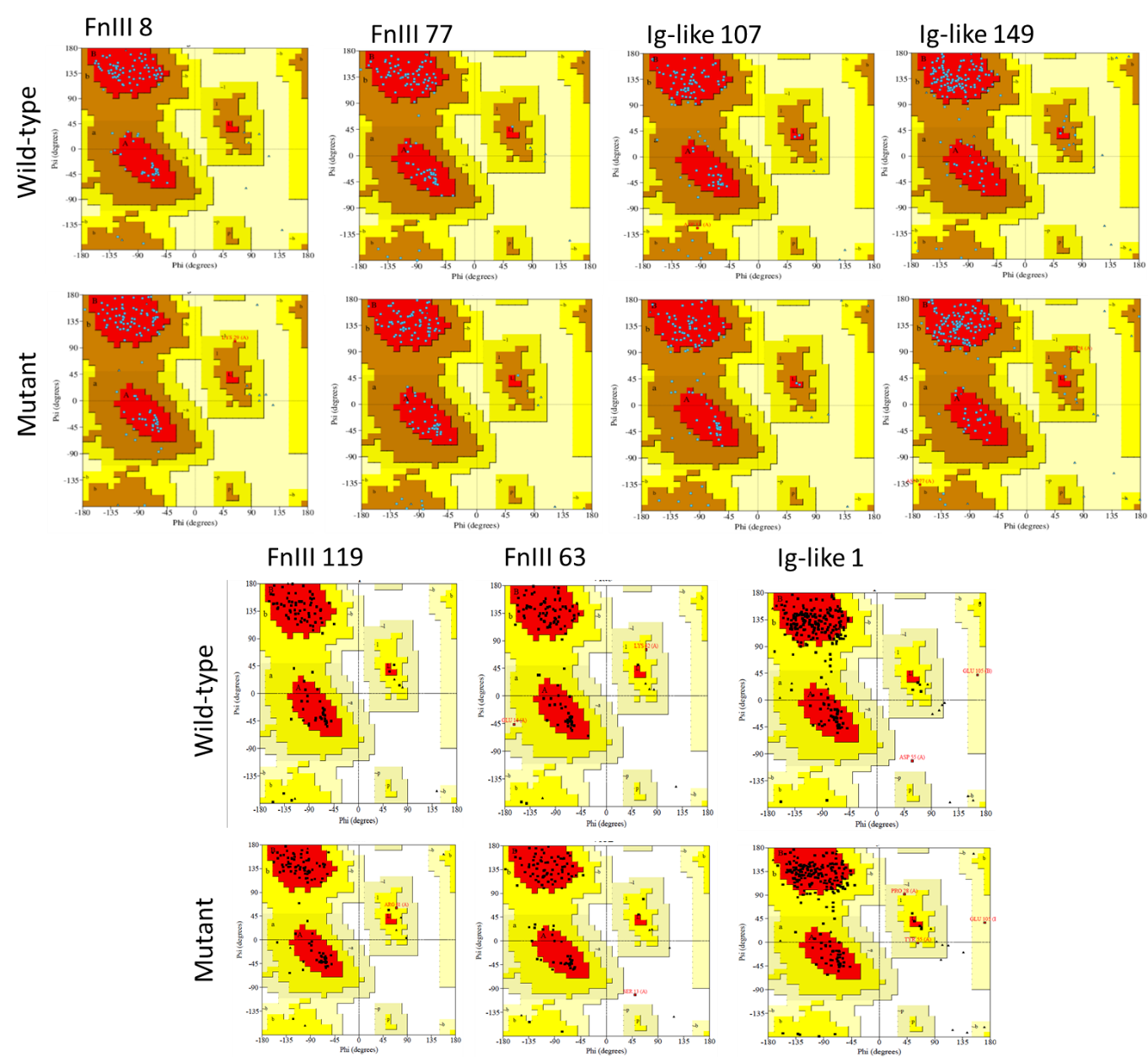

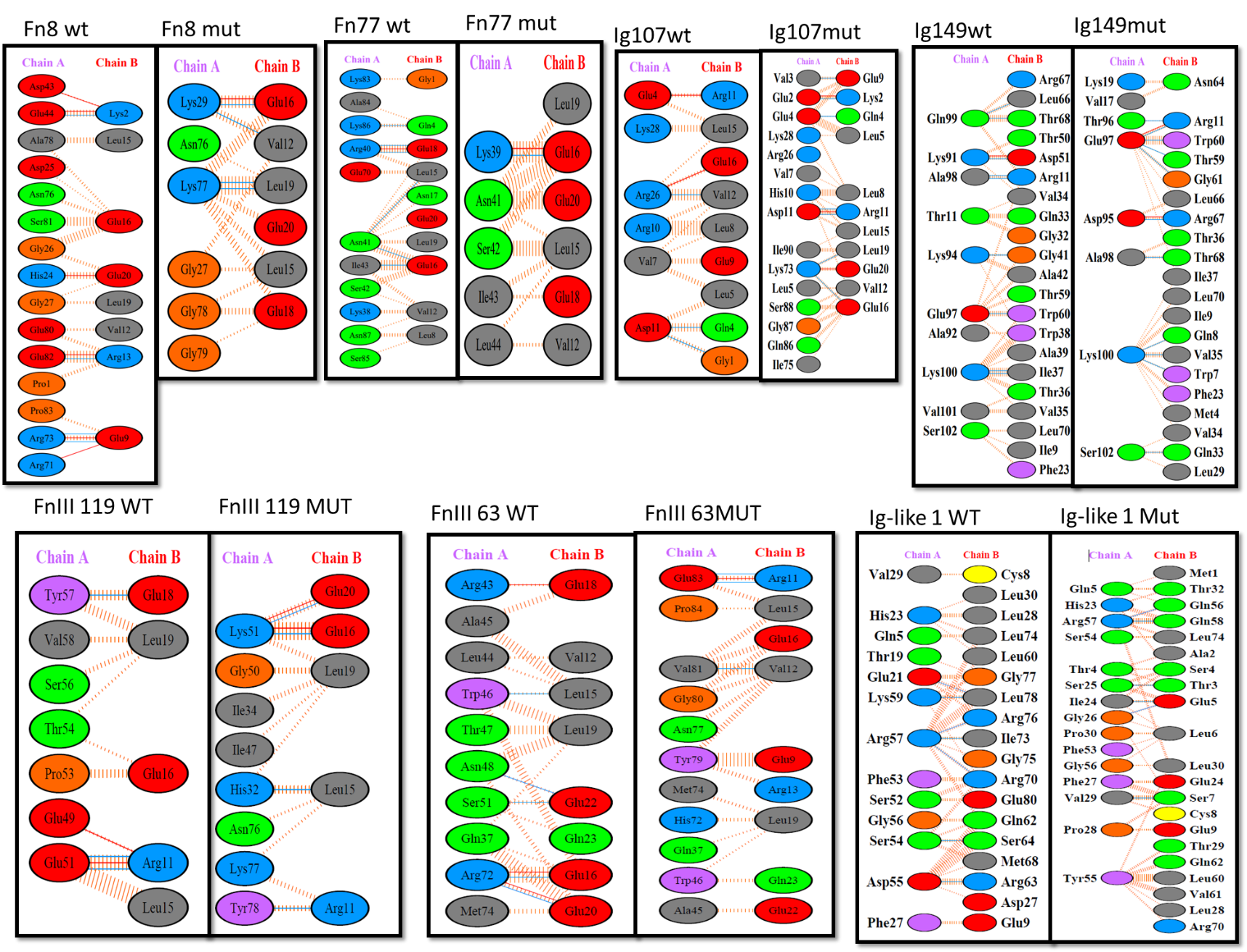
 ***Supplementary Figure 5: Representation of PDBsum data showing exact interacting residues of titin with its interacting partners; Chain A is titin and Chain B is TCAP (for Ig-like 1), Myosin (Fn8, Fn77, Ig 107, Fn63, Fn119) and Lamin (for Ig-like 149); Wild-type and mutant complex showing differences in their H-bonds, salt bridges and non-bonded interactions***

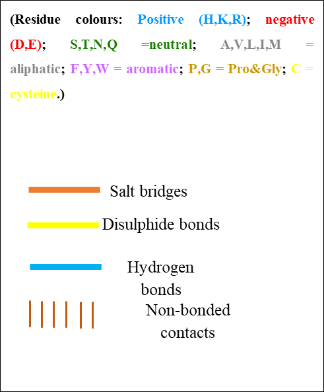
